# Essential roles for deubiquitination in *Leishmania* life cycle progression

**DOI:** 10.1101/2020.03.05.978528

**Authors:** Andreas Damianou, Rebecca Burge, Carolina M. C. Catta-Preta, Vincent Geoghegan, Y. Romina Nievas, Katherine Newling, Elaine Brown, Richard Burchmore, Boris Rodenko, Jeremy C. Mottram

## Abstract

The parasitic protozoan *Leishmania* requires proteasomal, autophagic and lysosomal proteolytic pathways to enact the extensive cellular remodelling that occurs during its life cycle. The proteasome is essential for parasite proliferation, yet little is known about the requirement for ubiquitination/deubiquitination processes in growth and differentiation. Activity-based protein profiling of *L. mexicana* C12, C19 and C65 deubiquitinating cysteine peptidases (DUBs) revealed DUB activity remains relatively constant during differentiation of procyclic promastigote to amastigote. However, when Bar-seq was applied to a pool of 16 DUB null mutants created in promastigotes using CRISPR-Cas9, significant loss of fitness was observed during differentiation and intracellular infection. DUBs 4, 7, and 13 are required for successful transformation from metacyclic promastigote to amastigote and DUBs 3, 5, 6, 10, 11 and 14 are required for normal amastigote proliferation in mice. DUBs 1, 2, 12 and 16 are essential for promastigote viability and the essential role of DUB2 in establishing infection was demonstrated using DiCre inducible gene deletion *in vitro* and *in vivo*. DUB2 is found in the nucleus and interacts with nuclear proteins associated with transcription/chromatin dynamics, mRNA splicing and mRNA capping. DUB2 has broad linkage specificity, cleaving all the di-ubiquitin chains except for Lys27 and Met1. Our study demonstrates the crucial role that DUBs play in differentiation and intracellular survival of *Leishmania* and that amastigotes are exquisitely sensitive to disruption of ubiquitination homeostasis.

**Author Summary:** *Leishmania* parasites require a variety of protein degradation pathways to enable the parasite to transition through the various life cycle stages that occur in its insect and mammalian hosts. Several enzymes involved in protein degradation in *Leishmania* are known to be essential, including a multi-protein complex, the proteasome, but little is known about how proteins are targeted to the proteasome for degradation. Here, we analyse components of the deubiqutination pathway, including twenty cysteine peptidases (DUBs) that remove the posttranslational modifier ubiquitin from substrates tagged for proteasomal degradation. We used chemical probes to measure active enzymes in parasite lysates and genome engineering to create DUB gene deletion mutants. We identified some DUBs that are essential for parasite viability and some that are required for life cycle progression. We carried out a detailed analysis of the essential DUB2, which has broad deubiquitinase activity and is found in the nucleus. This enzyme interacts with nuclear proteins associated with transcription/chromatin dynamics, mRNA splicing and mRNA capping. This work demonstrates the important role that DUBs play in *Leishmania in vivo* infection and further validates DUBs as potential drug targets in this parasite.

## Introduction

Leishmaniasis is a vector-borne neglected tropical disease that causes serious public health problems in 98 tropical and sub-tropical countries, and is responsible for 20,000 – 40,000 deaths annually [1]. The disease spectrum varies from asymptomatic infection, through a disfiguring and debilitating mucocutaneous form to a visceral form that can be fatal. The causative agent of this disease is the obligate intracellular parasite of the genus *Leishmania*. The developmental process of *Leishmania* is closely linked with the ability of this parasite to survive and replicate in distinct environmental niches. The medical importance of one of these environmental niches is evident in the phagolysosomes of the infected host macrophages, where the *Leishmania* parasites differentiate from the motile promastigote to the non-motile amastigote and subsequently proliferate and establishes an infection. The morphological and transcriptional changes that occur after differentiation of *Leishmania* promastigotes to amastigotes are well established [2], but little is known about the regulatory and operational processes that underpin this transition. Reversible post-translational modification (PTM) of proteins with chemical groups, such as phosphorylation, or other proteins, such as ubiquitination, are likely to be pivotal for successful life cycle progression [3–5] and intracellular parasitism. In addition, irreversible PTM such as proteolytic cleavage is important for the cellular remodeling that occurs during the life cycle of *Leishmania* and the commitment to differentiation [6–8], as well as in amastigote survival [9,10].

Ubiquitination is an evolutionarily conserved posttranslational modification involving a series of highly regulated enzymatic reactions resulting in attachment of ubiquitin to proteins in the cell [11]. Ubiquitination regulates vital cellular mechanisms including protein degradation through the ubiquitin-proteasome system (UPS), autophagy, DNA repair and protein trafficking. For the covalent attachment of ubiquitin to target protein or other ubiquitin, three types of enzymes are required to act sequentially: E1 ubiquitin-activating enzyme, E2 ubiquitin-conjugating enzyme and an E3 ubiquitin ligase [11]. This system is reversible and deubiquitinase enzymes (DUBs) are responsible for the removal of ubiquitin from the target protein, providing an additional level of regulation of the ubiquitination pathway [12]. The number of DUBs varies from organism to organism. For example, around 100 DUBs have been discovered in humans but only 20 DUBs in *Saccharomyces cerevisiae* [13]. In general, DUBs can be divided into seven distinct structural families [14]: ubiquitin-specific proteases (USPs, family C19), C-terminal hydrolyses (UCHs, family C12), ovarian tumor proteases (OTUs, family C65), JAB1/MPN/MOV34 metalloenzymes (JAMM/MPN+, family), Josephins, family C86, MINDY, family C115 [15] and the newly identified ZUFSP (zinc finger with UFM1-specific peptidase domain protein), family C78 [13,16]. Here we characterise DUBs of *Leishmania* and describe a Bar-seq CRISPR-Cas9 genome editing strategy that was used to evaluate the requirement for DUB activity in the *Leishmania* life cycle. Our data demonstrate that the broad and diverse mechanisms used by DUBs are required for successful *Leishmania* life cycle progression.

## Results

### *Leishmania* deubiquitinase cysteine peptidase gene family

We searched the genome of *L. mexicana* and identified 20 DUBs belonging to the C12, C19 and C65 families (Table S1). Nineteen of these 20 DUBs are conserved in the closely related trypanosamatid *Trypanosoma brucei*, with only the C19 family DUB18 being absent. Domain analysis using InterPro [17] confirmed the presence of the expected DUB peptidase domains. Furthermore, several other domains including UBA [18], zinc finger UBP-type [19], WD40, exonuclease RNase and PAN2 domains were also identified (Fig. 1a).

**Fig. 1.**
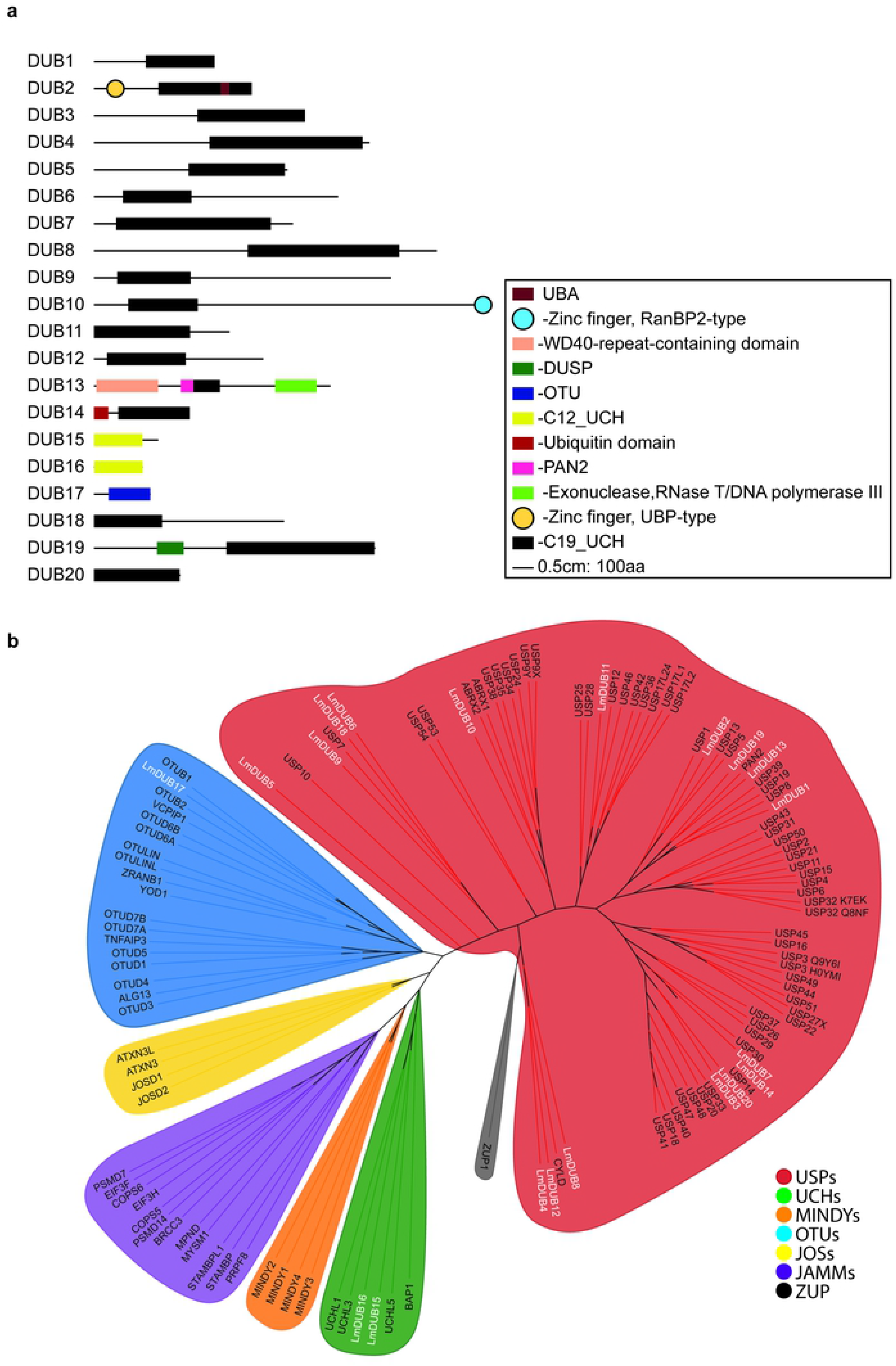
*Leishmania* DUBs domain analysis and phylogenetic tree. **a** Domain analysis of 20 predicted cysteine peptidase DUBs. The length of each line correlates to the protein size (see line scale). InterPro was used to analyse protein sequences. **b** Protein sequences for *Leishmania* and *H. sapiens* DUBs were retrieved from Uniprot and the DUB domain extracted and aligned using MUSCLE. Finally, a constraint phylogenetic tree was generated by RAxML. The constraint tree was created by including mammalian DUB domains into the seven known families, whereas *Leishmania* DUBs were free to cluster into any DUB family. The LG Substitution matrix was used. The best fit model tree was further designed initially in the iTOL INTERACTIVE TREE OF LIFE (https://itol.embl.de/) where branched length was ignored, and an unrooted tree formed.

A phylogenetic tree of *L. mexicana* and *Homo sapiens* DUBs revealed the clustering of *Leishmania* DUBs into human DUB familes. DUB15 and DUB16 clustered with mammalian C12 UCH DUBs and DUB17 with mammalian C65 OTU as expected (Fig. 1b). USP (C19) DUBs are the largest class of DUBs in humans (~60 members) and also in *Leishmania* (17 members). Most of the USPs include domains in addition to their C19 catalytic domain. The C19 DUBs have limited sequence conservation, restricted to the catalytic domain which consists of an active catalytic triad including Cys, His and Asp or Asn residues. The rest of the protein sequence is highly diverse among the USPs and it is believed that this sequence diversity is crucial for the substrate recognition through interaction with other proteins.

### *Leishmania* DUBs required for life cycle transition

The fluorescent activity-based probe (ABP) Cy5UbPRG, previously developed for its ability to react with active DUBs, identified active DUBs in the lysate of *L. mexicana* promastigotes [20]. In-gel fluorescence showed the presence of several DUBs in the range of 30-160 kDa (Fig. 2a), the size expected taking into account the covalent attachment of the 9 kDa Cy5UbPRG probe (Table S1). The activity of DUBs detected with the ABP was similar in promastigotes, axenic amastigotes and lesion-derived amastigotes and during differentation from amastigotes to promastigotes (Fig 2b). UbPRG was then attached to Sepharose beads and used to affinity purify proteins from an *L. mexicana* promastigote cell lysate, and mass spectrometry used to confirm DUB identity. Six DUBs (DUB2, DUB15, DUB16, DUB17, DUB18, and DUB19) were successfully identified only in the UbPRG sample and not in the control sample, indicating that the warhead can successfully react with cysteine peptidase DUBs (Fig. 2c and Table S2). Twenty other proteins specifically identified in the UbPRG sample are likely to be proteins that are DUB substrates (such as polyubiquitin, LmxM.09.0891), form complexes with DUBs, or react directly with the PRG warhead.

**Fig. 2.**
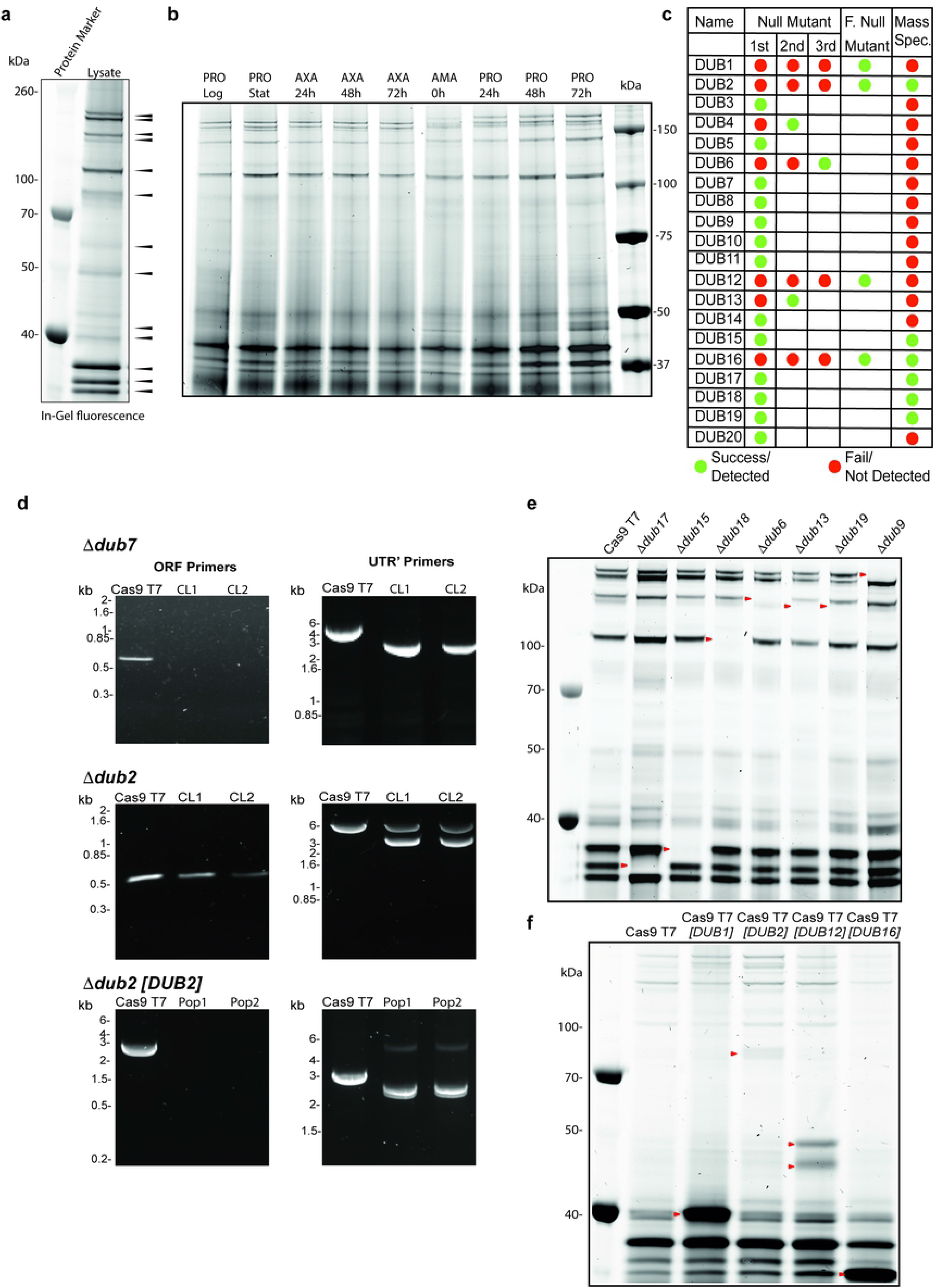
Validation of DUB null mutants. **a** DUB activity in *L. mexicana* promastigote cell lysates was profiled using Cy5UbPRG. Proteins were resolved by SDS-PAGE and in-gel fluorescence was captured using a Typhoon imager. Black arrows represent the higher intensity activities. **b** DUB activity profiled using Cy5UbPRG in log phase promastigotes (PRO Log), stationary phase promastigotes (PRO Stat), axenic amastigotes (AXA) grown for 24, 48 and 72 h after differentiation from stationary phase promastigotes, lesion-derived amastigotes (AMA) and promastigotes 24, 48 and 72 h after differentiation from AMA. **c** Summary of the generation of DUB null mutants and the detection of DUBs by affinity purification and mass spectrometry. Green, successful; red, not successful. F; facilitated **d** Example of diagnostic PCRs performed to confirm a null mutant for a non-essential gene (Δ*dub7*), a heterozygote for an essential gene (Δ*dub2*) and the generation of a facilitated null mutant (Δ*dub2 [DUB2]*) **e** DUB activity profiling of lysate extracted from null mutant lines of log-phase *L. mexicana* promastigotes. Arrow heads represent the absence of an active DUB compared to the Cas9 T7 parental cell line. **f** DUB activity profiling of lysate extracted from facilitated null mutant lines of log-phase *L. mexicana* promastigotes. The arrow heads represent the DUB expressed from the episome.

To assess whether the 20 DUBs were dispensable for growth of *L. mexicana* promastigotes each gene was targeted for deletion using CRISPR-Cas9 [21] with two barcoded antibiotic resistance cassettes used to attempt to replace both allelic copies of each *DUB* (Fig. 2c). In the first round of transfection into the parental *L. mex Cas9* T7 line, two independent null mutant lines were generated for 13 DUBs (Δ*dub3*, Δ*dub5*, Δ*dub7*, Δ*dub8*, Δ*dub9*, Δ*dub10*, Δ*dub11*, Δ*dub14*, Δ*dub15*, Δ*dub17*, Δ*dub18*, Δ*dub19*, Δ*dub20*) and these were validated using PCR to detect correct integration of the resistance cassettes and removal of the open reading frame (Fig. 2d, Fig. S1 and Table S3). After further transfections, another 3 null mutants were isolated (Δ*dub4*, Δ*dub6* and Δ*dub13*) giving a total of 16. For four DUBs, *DUB1*, *DUB2*, *DUB12*, *DUB16*, null mutants could not be generated.

To identify the DUB activities detected in promastigotes (Fig. 2a), cell lysates for the *DUB* null mutants were probed with the Cy5UbPRG ABP (Fig. 2e, Fig S2.), with the expectation that the absence of a band of activity in comparison to the *L. mexicana Cas9* T7 would represent the DUB that had been deleted. This allowed us to identify 7 active DUBs close to their expected sizes; DUB6 (133 kDa), DUB9 (155 kDa), DUB13 (126 kDa), DUB15 (34 kDa), DUB17 (30 kDa), DUB18 (102 kDa) and DUB19 (126kDa). The fact that DUB15, DUB16, DUB17, DUB18, and DUB19 had been affinity purified with the UbPRG probe shows that these DUBs are active in the promastigote stage. The same sized DUB activity was observed to be absent in both Δ*dub13* and Δ*dub19;* the estimated size suggests this might be DUB13 (Fig. S2). Several null mutant DUB cell lines did not show any depletion of activity with the CY5UbPRG profiling, suggesting that those DUBs are not expressed, not active or are below detection limits of the ABP in the promastigote stage under the environmental conditions tested or that the DUBs do not bind the ABP. To investigate further, cell lines were differentiated to amastigote and probed with the ABP. This allowed DUB8 (172kDa) and DUB19 (148kDa) to be identified by comparing wild type and Δ*dub8* and Δ*dub19* (Fig. S3b).

It was not possible to isolate null mutant clones for *DUB1*, *DUB2*, *DUB12* and *DUB16*, possibly because those DUBs are essential in promastigotes (Fig. 2c). In three instances (*DUB1*, *DUB2* and *DUB12*), correct integration of the drug resistance cassettes was observed, but the parasites retained a wild type copy of the gene, indicating that the genes are essential (Fig. S1). For *DUB16*, no populations could be recovered after the transfections. To provide further evidence that the genes are essential and to discount the possibility that the lack of gene deletion was a technical failure, facilitated null mutants were generated for the four *DUB* genes. Initially, the parental Cas9 T7 cell line was transfected with a plasmid expressing the DUB, thereafter the two alleles in the genomic locus were deleted successfully using CRISPR, two independent populations isolated, and the expected gene editing confirmed by diagnostic PCR (Fig. 2d and Fig. S1). The Cy5UbPRG ABP was used to identify the over-expressed and active DUBs in the *L. mexicana* Cas9 T7 [*DUB*] cell lines (Fig. 2f), where an increase in specific activities were detected in the over-expression lines compared to the parental line (Fig. 2f, arrow heads). For DUB2 and DUB16 the expected sizes of 81 and 25 kDa were detected respectively, once the ~9 kDa mass of the ABP had been taken into account. In contrast, both DUB1 (expected size 65 kDa, observed size 40 kDa) and DUB12 (expected size 89 kDa, observed sizes 45 and 50 kDa) are smaller than expected and seem likely to be proteolytically processed in the cell (Fig. 2f).

### Bar-seq reveals the importance of DUBs in the *L. mexicana* life cycle

To assess the ability of the DUB null mutants to transition through the parasite life cycle, Bar-seq analysis was carried out on pools of barcoded null mutants [22]. The 58 null mutants representing 16 DUBs, 13 other peptidases, 25 other ubiquitination system genes and 4 protein kinases (Table S4) were prepared in equal proportions in 6 replicate promastigote (PRO) pools. The pools were allowed to grow to stationary phase before samples were collected and either induced to form axenic amastigotes (AXA) or used to purify metacyclic stage promastigotes (META). Purified metacyclic promastigotes were used to infect macrophages (inMAC) *in vitro* or mice via footpad (FP) injection. At various time points, DNA samples were prepared from the pool for amplification of the barcodes by PCR and quantitative analysis via next generation sequencing (Fig. 3a, Table S4).

**Fig. 3:**
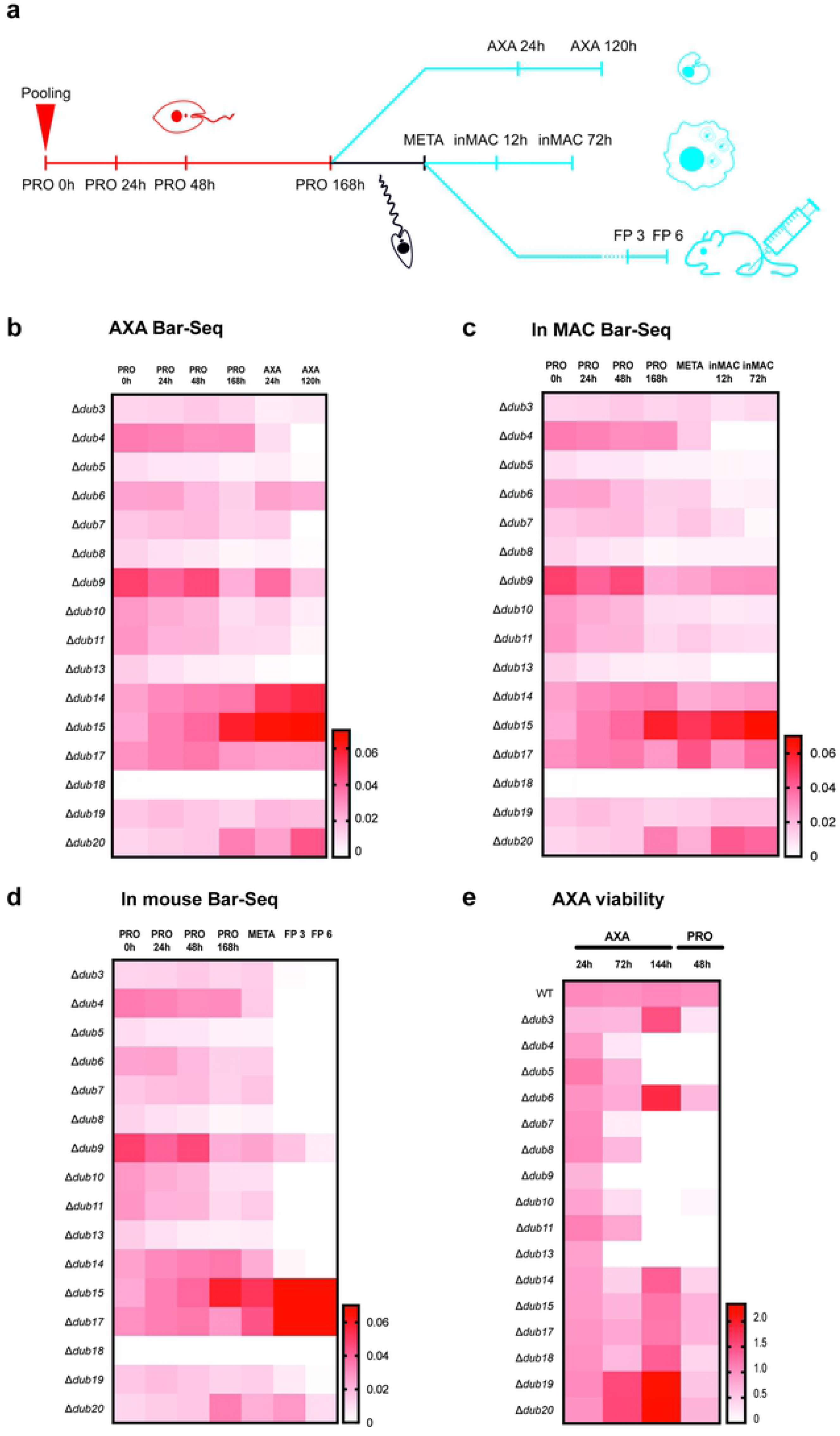
Life cycle phenotyping of DUB null mutants by Bar-seq. Fifty-eight null mutants were pooled (n = 6) as promastigotes and grown to stationary phase before being induced to differentiate to axenic amastigotes *in vitro* or infect macrophages or mice. At the time points indicated in **a**, DNA samples were extracted for barcode amplification by PCR and quantitative analysis by next generation sequencing. Average proportional representation of null mutant-specific barcodes at each experimental stage is displayed in the heat maps for **b** axenic amastigote, **c** macrophage infection and **d** mouse infection experiments. Samples included represent promastigote time-point zero (PRO 0h), early-log phase (PRO 24h), mid-log phase (PRO 48h), late-log phase (PRO 72h), stationary phase (PRO 168h), early axenic amastigote differentiation (AXA 24h), post-axenic amastigote differentiation (AXA 120h), purified metacyclics (META), early macrophage infection (inMAC 12h), late macrophage infection (inMAC72h), 3 week footpad mouse infection (FP 3) and 6 week footpad mouse infection (FP 6). Separately, individual null mutant lines were induced to differentiate into amastigotes and cell viability measured at the stages indicated in **e** using AlamarBlue reagent (added 24 h prior to each measurement). A transformation step back to promastigotes was included. Measurements of cell viability were calculated relative to the wild-type at each experimental stage. The data are an average of two independent experiments.

In order to analyse fitness phenotypes, the number of reads for each barcode at each experimental stage were first divided by the total number of reads at the relevant stages. Next, data was averaged across the 6 replicates and the barcodes matched back to the original null mutant lines. Changes in the average proportional representation of these lines were then used to infer changes in fitness. Only small changes in fitness were observed between adjacent promastigote stages (PRO 0 h-PRO 168 h) but strong fitness defects were common upon entering the amastigote life cycles stages (AXA 24 h onwards, inMAC 12 h onwards and 3 Weeks FP onwards, Fig. 3b-d respectively). Across the three experiments, Δ*dub4*, Δ*dub7* and *Δdub13* had the strongest and most consistent loss of fitness phenotypes. Interestingly, although the data for the axenic amastigote and macrophage Bar-seq experiments is highly consistent, many additional loss-of-fitness defects were observed during mouse infection. Indeed, Δ*dub3*, Δ*dub5*, Δ*dub6*, Δ*dub8*, Δ*dub10*, Δ*dub11* and Δ*dub14* exhibited strong loss-of-fitness during mouse infection, despite not doing so in the *in vitro* experiments. While significant loss of fitness in mouse infection was observed for DUBs and other ubiquitination system mutants studied (Table S4), loss of fitness phenotypes were less common for other peptidase mutants in the screen, including members of the calpain (C2), SUMO-specific peptidase (C97) and D-alanyl-glycyl endopeptidase (C51) families [23].

To demonstrate the validity of the phenotypes observed in the Bar-seq screen, null mutants were also analysed individually at 24, 72 and 144 h after induction of amastigote differentiation and 48 h after transformation of amastigotes back to promastigotes using resazurin, an oxidation-reduction indicator that reports on cell activity. The data from this experiment matches well with the phenotypes observed in the AXA Bar-seq data (Fig. 3e). An exception to this is Δ*dub9* which is well represented throughout the Bar-seq experiment but loses viability at 72 h in the cellular assay, perhaps indicating the persistence of Δ*dub9* beyond 72 h but in a metabolically inactive state. Interestingly, Δ*dub18* survives well in the viability assay, showing that the relatively low representation of Δ*dub18* at the 0 h time point probably accounts for its low abundance throughout the Bar-seq experiments. The viability data also supports more unusual phenotypes. For example, Δ*dub6* is still well represented at the AXA 24 h and 120 h stages of the Bar-seq experiment but not at the in MAC 12 h stage. This is supported by data from the viability assay showing that Δ*dub6* is viable throughout amastigote differentiation. Based on this finding, one could consider that DUB6 may be specifically required for part of the infection process such as parasite entry into macrophages or survival in the phagolysosomes. Overall, the phenotypes observed in the pooled context of the Bar-seq experiment reflect well the individual phenotyping of the viability assay. To further validate how these null mutants behave during amastigote differentiation, a western blot against HASPB, a marker for amastigotes [24], was performed. Δ*dub4*, Δ*dub7*, Δ*dub9* and Δ*dub13* showed very low or no expression of HASPB 120 h after the initiation of differentiation (Fig. S3). These results demonstrate the inability of Δ*dub4*, Δ*dub7*, Δ*dub9*, and Δ*dub13* to differentiate into amastigotes.

### Location of DUBs in *L. mexicana*

To determine the cellular localisation of DUBs, a CRISPR-Cas9 approach [21] was used to endogenously tag the proteins at the N-terminus with mNeonGreen (mNeo). The use of N-terminal tagging was selected as most DUBs have their catalytic domain close to the C-terminus. All DUBs were successfully tagged except DUB3, for which no populations were isolated after either N- or C-terminal tagging attempts. Successful integration of the mNeo protein was confirmed using the Cy5UbPRG ABP (Fig. S2). The fusion of mNeo adds an extra 27 kDa to the predicted size of the DUB (Table S1) and a fusion protein of the expected size was detected for DUB2, DUB6, DUB9, DUB15, DUB16, DUB17, DUB18 and DUB19 (Fig. S2c), but not for the remaining DUBs. In some cases, such as DUB2 and DUB17, the native DUB could no longer be detected, suggesting that both alleles had been tagged. Nevertheless, several lines such as DUB16 and DUB18 expressed both native and tagged DUBs showing that only one allele was tagged. DUB4, DUB5, DUB8, DUB14 and DUB20 could not be identified either as fusion proteins, or by comparing wild type and knockout activity profiles, indicating that the proteins may be expressed at low levels, inactive or unable to bind the ABP (Fig. S2). Active DUB1 appears to be processed in the promastigote stage, so the mNeonGreen fusion may be cleaved. This is supported by the weak cytoplasmic fluorescence observed (Fig. S4), likely a result of the expression of the precursor protein. On the other hand, the DUB12::mNeon cell line had a strong nuclear signal, even though the full length protein could not be detected. DUB12 is essential for the survival of promastigotes and the detection of two processed and active forms of DUB12 (Fig. 2e), suggests that inactive full length DUB12 is trafficked to the nucleus where it is cleaved and activated. Cellular landmark analysis [25] revealed the localisation of DUBs to the cytoplasm (DUB1, DUB6, DUB8, DUB9, DUB10, DUB13, DUB15, DUB17, DUB18, DUB19, DUB20), nucleus (DUB2, DUB4, DUB5, DUB12, DUB14) and cytoplasm/flagellum (DUB16) (Fig.S4). Additionally, there were some DUBs with cellular landmarks [25] consistent with localisation to the mitochondrion (DUB11), endoplasmic reticulum (DUB7), lysosome (DUB2) and basal body (DUB6).

### DUB2 activity is essential for *L. mexicana*

*L. mexicana* DUB2 is one of the four essential DUBs identified in the CRISPR-Cas9 screen and is classified as a member of the USP family, closely related to mammalian USP5, USP13 and USP1 (Fig. 1b). DUB2 domain analysis suggested the presence of a peptidase C19, ubiquitin carboxyl-terminal hydrolase domain, a UBA (ubiquitin-associated domain and a Zinc finger UBP-type (Znf_UBP)) (Fig. 1a). The UBA domain found in the C-terminal end of DUB2 is present in many proteins associated with the ubiquitin system, and it is responsible for the incorporation of ubiquitinated substrates [26]. The Zinc finger domain is found only in a minor subfamily of USP DUBs including the mammalian USP5, USP13 and USP39, regulators of deubiquitination [19].

A modified version of the inducible DiCre system [27] was applied to *L. mexicana DUB2* to investigate its function in promastigotes. The first allele of *DUB2* was replaced with a floxed-*DUB2* (Δ*dub2::DUB2*^FLOX^/*DUB2*, called DUB2^+/+FLOX^). Subsequently, the second allele of *DUB2* was replaced with a *HYG* resistance cassette (Δ*dub2*::*HYG*/Δ*dub2*::*DUB2*^FLOX^,called *DUB2*^−/+FLOX^). The expected genetic modifications for several clones were confirmed by PCR (Fig. S5). Excision of *DUB2* in *DUB2*^+/+FLOX^ and *DUB2*^−/+FLOX^ logarithmic phase promastigotes was confirmed by the PCR amplification of a 1.3 kb DNA fragment in only those cells treated with 100 nM rapamycin (Fig. 4a). A pronounced defect in cell growth was observed in the rapamycin treated DUB2^−/+FLOX^ cell line in comparison to DUB2^+/+FLOX^ or untreated controls, indicating that deletion of both *DUB2* alleles is required to elicit a growth phenotype (Fig. 4b). The Cy5UbPRG ABP demonstrated that the activity of DUB2 sharply decreased in induced DUB2^−/+FLOX^ line, but not in DUB2^+/+FLOX^ or WT (Fig. 4c). For further analysis, the in-gel fluorescence band intensity was quantified for the DUB2^−/+FLOX^ line, showing one band (band 9) had a 70% depletion after rapamycin induction Fig. 4d). Band 9 is the same size as that observed in the episomal over-expression of DUB2 (Fig. 2e) and correlates with the predicted mass of DUB2 protein covalently bound to the ABP (~100 kDa). A second unidentified DUB, labelled Unk 4, had a slight increase in activity after depletion of DUB2 in the DUB2^−/+flox^ line (Fig. 4c,d). Loss of DUB2 may cause a compensatory change in activity for this DUB.

**Fig. 4:**
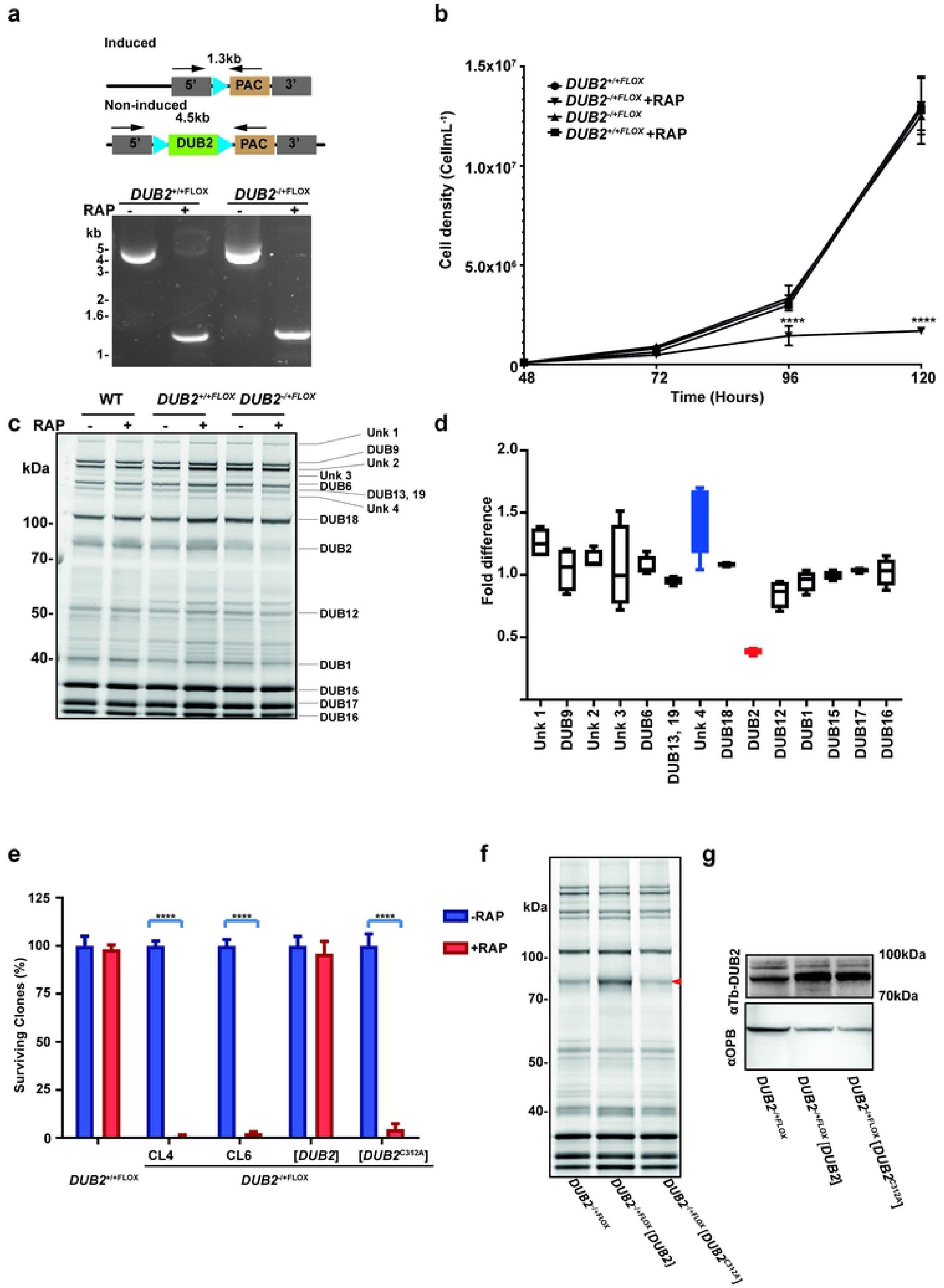
Conditional deletion of *DUB2*. **a** (Top) Schematic representation of the loxP-flanked *DUB2* allele before and after rapamycin-induced recombination by DiCre. Arrows represent the primers used for the PCR to check for the successful excision of *DUB2*. (Bottom) PCR amplification of *DUB2^−+/+FLOX^* and *DUB2^−/+FLOX^* +/- 100 nM RAP for 48 hours was conducted and the resulting amplicons resolved on 1% agarose gel stain with SYBR safe. **b** *DUB2^+/+FLOX^* and *DUB2^−/+FLOX^* promastigotes were seeded at a density of 1 × 10^5^ cells mL^−1^ and grown in the absence or presence (-/+) of 100 nM RAP for 48 h. Cells were diluted to 1 × 10^5^ cells mL^−1^ and allowed to grow in the presence and absence of 100 nM RAP for 4 d. Cell density was determined by counting at 24 h intervals and mean ± SD of triplicate values were plotted. The data were analysed in Prism software, and a t-test performed to indicate the significance (n=5). Error bars show the standard deviation of the mean. ****p < 0.0001. **c** DUBs visualised by in-gel fluorescence analysis after separation by SDS PAGE. Samples were lysed, and 25 μg of lysate was incubated with the probe for 30 min. The positions of the different DUBs are indicated. **d** Fold difference of the DUB activity comparing *DUB2* induced/*DUB2* non-induced, data from **c**. Band intensities were quantified by densitometry using GelAnalyzer 19.1 software. Each band is represented as a percentage of the total activity. The percentages for each band for the induced sample were divided against the non-induced to determine the fold difference. The data were then analysed in Prism software. Each column represents the mean of data from three independent experiments. Error bars show the standard deviation of the mean. Blue represents an increase of more than 1.3-fold whereas red represents a difference of less than 0.7-fold. **e** Percentage of surviving clones of *DUB2^+/+FLOX^*, *DUB2^−/+FLOX^* clone 4, *DUB2^−/+FLOX^* clone 6, *DUB2^−/+FLOX^* [*DUB2*] and *DUB2^−/+FLOX^* [*DUB2*^C312A^] promastigotes after RAP induction. The cell lines were incubated with or without RAP for 48 hours. Then, cells were counted and cloned into 96 well plates. After 4 weeks plates were checked for surviving parasites. The data were then collected and analysed in Prism software. 100% represents the highest number of surviving parasites that were collected from an individual experiment in the absence of RAP. Each column represents the mean of data from three independent experiments. Error bars show the standard deviation of the mean. ****p < 0.0001. **f** DUBs were visualised by in gel fluorescence analysis after separation by SDS-PAGE. Samples were lysed and 25 μg of lysate was incubated with the probe for 30 min. Lysate was collected from *DUB2^−/+FLOX^*, *DUB2^−/+FLOX^* [DUB2] and *DUB2^−/+FLOX^* [DUB2^C312A^] promastigotes in the presence of 10 μg/μL of G418. **g** Western blotting analysis with anti-TbDUB2 (1;1,000) and anti-LmOPB (1:20,000) (Loading control) antibodies on protein extracted from *DUB2^−/+FLOX^*, *DUB2^−/+FLOX^* [DUB2] and *DUB2^−/+FLOX^* [DUB2^C312A^] promastigotes grown in the presence of 10 μg/μL of G418.

To confirm that the depletion of DUB2 leads to cell death a clonogenic assay was performed on *DUB2*^+/+FLOX^ and *DUB2*^−/+FLOX^ promastigotes following treatment with rapamycin or DMSO for 48 h. A >90% decrease in numbers of clones surviving for both *DUB2*^−/+FLOX^ clones was observed for cells treated with rapamycin, whilst there was no difference in the number of clones surviving in the *DUB2*^+/+FLOX^ line, despite the presence of rapamycin (Fig. 4e). These data confirm that the loss of *DUB2* leads to death of promastigotes and that one allele of *DUB2* can sustain growth. Finally, diagnostic PCR confirmed the presence of *DUB2* in *DUB2*^−/+FLOX^ cells that survived rapamycin treatment, showing that if *DUB2* is deleted, the cells were non-viable (not shown).

Alignment of the C19 domain of DUB2 with other DUBs allowed the identification of C312 as the likely active site cysteine. A point mutation was introduced into *DUB2* in a pNUS plasmid to change the encoded cysteine 312 to alanine, which would result in abolishment of DUB2 activity. pNUS-*DUB2* and pNUS-*DUB2*^C312A^ plasmids were transfected into *DUB2*^−/+FLOX^ and an ABP assay with Cy5UbPRG used to confirm an increase of DUB2 activity in *DUB2^−/+FLOX^* [DUB2] but not *DUB2^−/+FLOX^* [DUB2^C312A^] (Fig. 4f). Over-expression of DUB2 was confirmed in both *DUB2^−/+FLOX^* [DUB2] and *DUB2^−/+FLOX^* [DUB2^C312A^] cell lines by western blot using an antibody raised against *T. brucei* DUB2; this cross-reacts with *L. mexicana* DUB2 (Fig. 4g). These data confirm that C312 is the active site cysteine and is required for DUB2 activity. The loss of the *DUB2* following rapamycin treatment resulted in a 90% decrease in the number of *DUB2^−/+FLOX^* [DUB2^C312A^] clones in comparison to *DUB2^−/+FLOX^* [DUB2] (Fig. 4e); indeed, expression of DUB2 in the *DUB2^−/+FLOX^* line resulted in full complementation of the cell death phenotype. Taken together, these experiments strongly support that the deubiquitinase activity of DUB2 is required for the survival of *L. mexicana* promastigotes.

The *DUB2*^−/+FLOX^ line was also used to examine if *DUB2* is essential for infection and survival in BALB/c mice. Mid-log phase WT and *DUB2*^−/+FLOX^ were grown in the presence or absence of rapamycin for 72 h and then inoculated into BALB/c mice. Efficient excision of DUB2 was confirmed by PCR (Fig. 5a). The mice infected with the rapamycin-treated WT or *DUB2*^−/+FLOX^ showed a steady growth in the footpad size whereas rapamycin-treated mice *DUB2*^−/+FLOX^ did not develop a footpad lesion (Fig. 5b), had 100-fold less parasites than non-treated in the footpad and could not be detected in the popliteal draining lymph nodes (Fig 5c). The presence of a small number of parasites in the footpad of treated mice might be due to the presence of a subpopulation of cells that retained the floxed *DUB2*. This was supported by the finding that DUB2 could be detected by PCR in cells isolated from treated *DUB2*^−/+FLOX^ lesions (Fig 5d). In summary, *DUB2* is required either for the establishment and/or maintenance of an infection in mice.

**Fig. 5:**
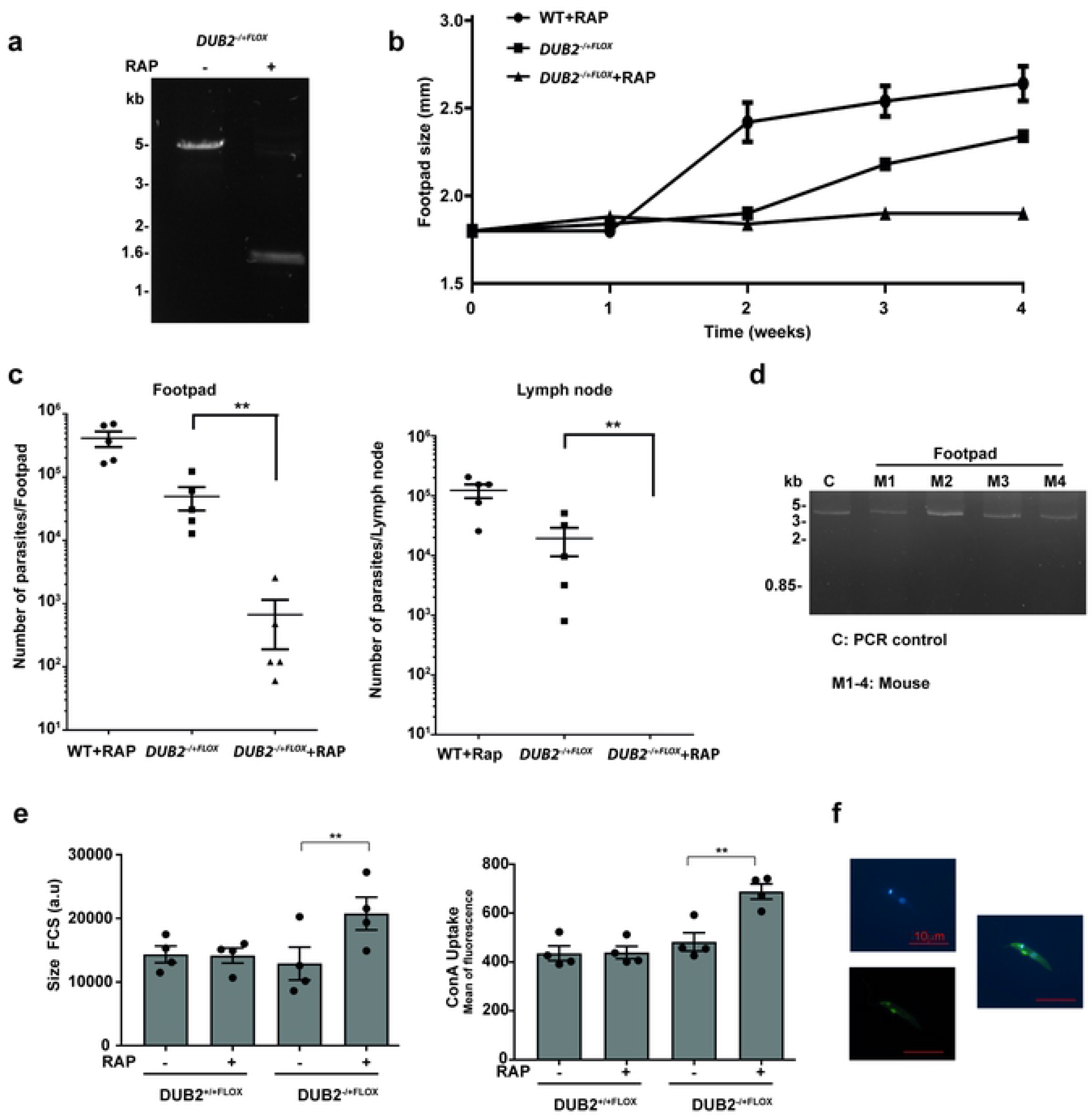
In vivo characterisation of *DUB2*. **a** PCR amplification of the floxed *DUB2* locus of *DUB2^−/+FLOX^* of 4 × 10^6^ cells mL^−1^ log phase promastigotes after incubation in the presence (+) or absence (-) of 1 μM RAP for 72 h. The resulting amplicons were resolved in 1% agarose gel stain by SYBR safe. Schematic representation of the primers used for the PCR, as well as the expected fragments (4.5kb, 1.3kb), are shown in Fig. 4a. **b** Mice were infected in the footpad with 2 × 10^6^ WT+RAP, DUB2^−/+FLOX^ and DUB2^−/+FLOX^+RAP promastigotes and footpad size was recorded by weekly caliper measurement. Data were analysed in Prism software. Data shown represent the mean footpad size and SD from groups of five mice. **c** Parasite burden. Mice were sacrificed after 4 weeks, and parasite loads in footpad and popliteal lymph node were determined by limiting dilution. Statistical analyses were performed in Prism software using the paired t-test. **d** PCR amplification to check *DUB2* excision of the samples collected from the mice which were infected with the *DUB2^−/+FLOX^* +RAP. The resulting amplicons were resolved in 1% agarose gel stained with SYBR safe. Schematic representation of the primers used for the PCR, as well as the expected fragments (5.6 kb, 1.3 kb), are shown in Fig. 4a. **e** Compiled flow cytometry results of the forward scatter (Left) or ConA uptake (Right) in DUB2^+/+FLOX^ and DUB2^−/+FLOX^ promastigotes Clone 4 and Clone 6 after 24 hours post induction. Cells were grown for 48 h with or without rapamycin. Afterwards, the cells were seeded at a density of 1 × 10^5^ cells mL^−1^ and allowed to grow in the presence or absence of RAP for 24 hours. Figure was generated in Prism Software. Each band represents a mean of 4 technical replicates and 2 biological replicates. Finally, a paired t-test was performed to assess statistical difference between non-induced and induced samples. Error bars represent SEM. **f** Cells were visualised under the microscope to check the correct localisation of ConA. Imaging of parasites was performed using confocal microscopy and processed in the ZEN 2.3 software. Panel contents: top left, DAPI staining; bottom left, the green fluorescence signal from ConA-FITC green; right, merge.

The likelihood that depletion of a DUB triggers secondary effects due to reduction of the free ubiquitin pool makes it challenging to characterise the function of a single DUB. Regulation of endocytosis by DUBs is well defined in mammalian cells [28], and one of the most fundamental processes in *Leishmania* is endocytosis, which is responsible for acquisition of essential molecules from the environment. In *Leishmania* endo- and exocytosis takes place in the flagellar pocket and uptake of fluorescently labelled Concanavalin A (ConA) was used to monitor any defects in the rapamycin-induced *DUB2*^+/+FLOX^ and DUB2^−/+FLOX^ mutants (Fig. 5e, f). After induction, DUB2^−/+FLOX^ mutants were bigger, as determined by forward scatter FSC-1 (Fig. 5e Left), and had increased endocytosis (Fig. 5e Right and Fig. 5f).

### The DUB2 interactome

To gain more insight into DUB2 function, the DUB2 interactome was investigated. The expression of active N-terminally tagged 3 × myc mNeo DUB2 was confirmed with the ABP and Western blotting (Fig. S6). DUB2 and an N-terminally tagged 3 × myc mNeo RDK2 control were immune-precipitated from *L. mexicana* promastigote cell lysates and proteins identified by mass spectrometry. Parasites were cross-linked to enable transient interactors to be captured. Data were processed with an interaction scoring algorithm SAINTq to provide a score on the probability of a true interaction with DUB2. This analysis produced a list of 110 high confidence DUB2 interactors at <1% FDR (Fig. 6, Table S5 that includes a number of proteins involved in ubiquitination, including poly-ubiquitin and three putative E2 ubiquitin-conjugating enzymes (*LmxM.34.1300*, *LmxM.04.0680*, *LmxM.13.1580*). Among the most enriched proteins was CYC12 (*LmxM.36.5640*), an L-type cyclin involved in spliced leader trans-splicing of pre-mRNA [29]. Other proteins involved in mRNA splicing were also enriched including SmD2, U5 snRNP and snRNP-B. Three members of the cleavage and polyadenylation specificity factor complex (CPSF) and a cleavage stimulation factor co-purified, suggesting a role for DUB2 in 3’ processing of mRNA. Proteins involved in mRNA capping, CBP110 and CGM1, were also strongly enriched with DUB2. In addition, 19 of the interactors are either annotated as RNA binding or known to have a role in RNA metabolism. A smaller number of DUB2 interactors are implicated in transcription, however, this group includes the top enriched interactor NIF3. Interestingly, among this group of proteins involved in transcription/chromatin dynamics is Nucleosome Assembly Protein, NAP, which was enriched 27-fold. In humans, NAP1 is a histone chaperone and plays an important role in nucleosome assembly/disassembly by binding to free H2A/H2B dimers [30]. Cycles of ubiquitination and de-ubiquitination of H2B are well known to play an important role in nucleosome dynamics during transcription and DNA replication. Several proteins with key roles in DNA replication were also identified as DUB2 interactors, such as PCNA, DNA polymerase delta and subunits of replication factor A and C. In other organisms, multiple components of the DNA replication machinery are ubiquitinated, in particular, ubiquitination of PCNA is critical for recruitment of DNA polymerases [31]. The identification of VPS4 [32] and a dynamin-1 like protein suggest involvement of DUB2 in endosomal trafficking. Finally, 3 protein kinases were co-enriched, including a mitogen activated protein kinase kinase (MKK1). This suggests a mechanism of DUB2 regulation, therefore mass spectrometry data were searched for phosphorylation sites revealing that DUB2 is phosphorylated on a serine in the region 586-589. A ubiquitination site in DUB2, K681, was also identified suggesting that DUB2 itself may be regulated by the ubiquitination pathway (Table S6). Overall, the multiple interactions of DUB2 with splicing factors, proteins involved in mRNA capping and factors regulating chromatin dynamics/transcription, place DUB2 near actively transcribing genes.

**Fig. 6:**
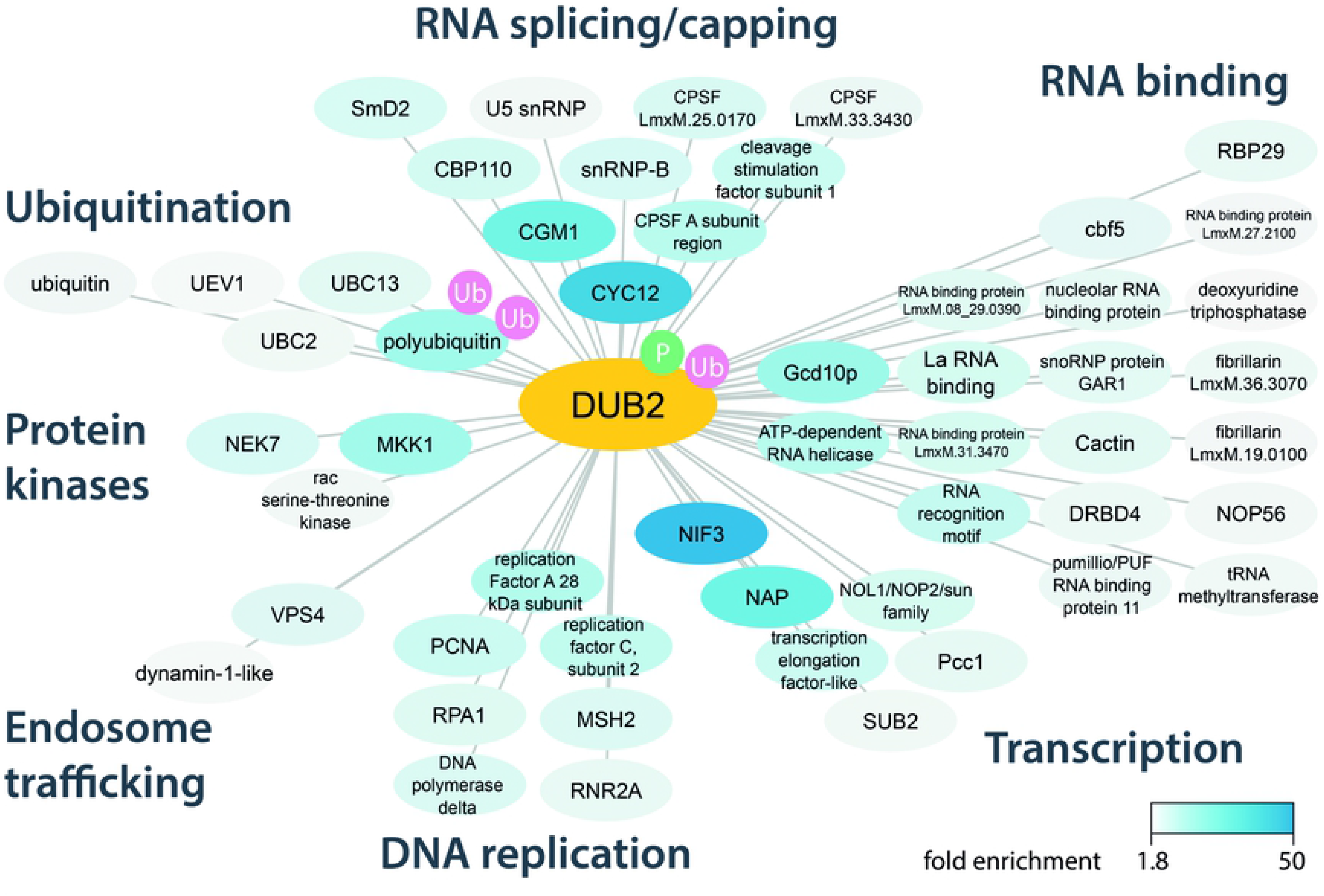
DUB2 interactome. Summary of detected DUB2 interactions. Myc-DUB2 was immuno-precipitated from promastigotes and binding partners identified by label free quantitative mass spectrometry. Interaction data were analysed with SAINTq, high confidence interactors (<1% FDR) were selected and are shaded according to fold enrichment against a control immuno-precipitation. Detected ubiquitination sites (Ub) and phosphorylation (P) are shown. Interactors are grouped according to annotated function. A total of 110 high confidence interactors were identified, non-annotated or ribosomal proteins have been omitted for clarity.

### DUB2 has deubiquitinating activity

To explore the function of DUB2 in more detail, recombinant enzyme (DUB2) and a catalytically inactive form (DUB2^C312A^) was expressed and purified using a baculovirus insect cell expression system (Fig. S7). DUB2, but not DUB2^C312A^ recombinant protein reacted with the Cy5UbPRG probe (Fig. 7a) and 85% of the recombinant DUB2 reacted with the probe (Fig. 7b). DUB2, but not DUB2^C312A^ recombinant protein had activity towards Ub-AMC substrate with a k_cat_ of 0.049 s^−1^ and a K_m_ of 14.9 × 10^−6^ M. DUB2 linkage specificity against all eight types of di-ubiquitin was then investigated over 30 min at a constant enzyme and substrate concentration. At 0.02 nM DUB2 had a preference for cleavage at Lys-11, Lys-29, Lys-33 and Lys-63, but at 2 nM DUB2 cleaved all linkages except Lys-27 (Fig. 7c).

**Fig. 7:**
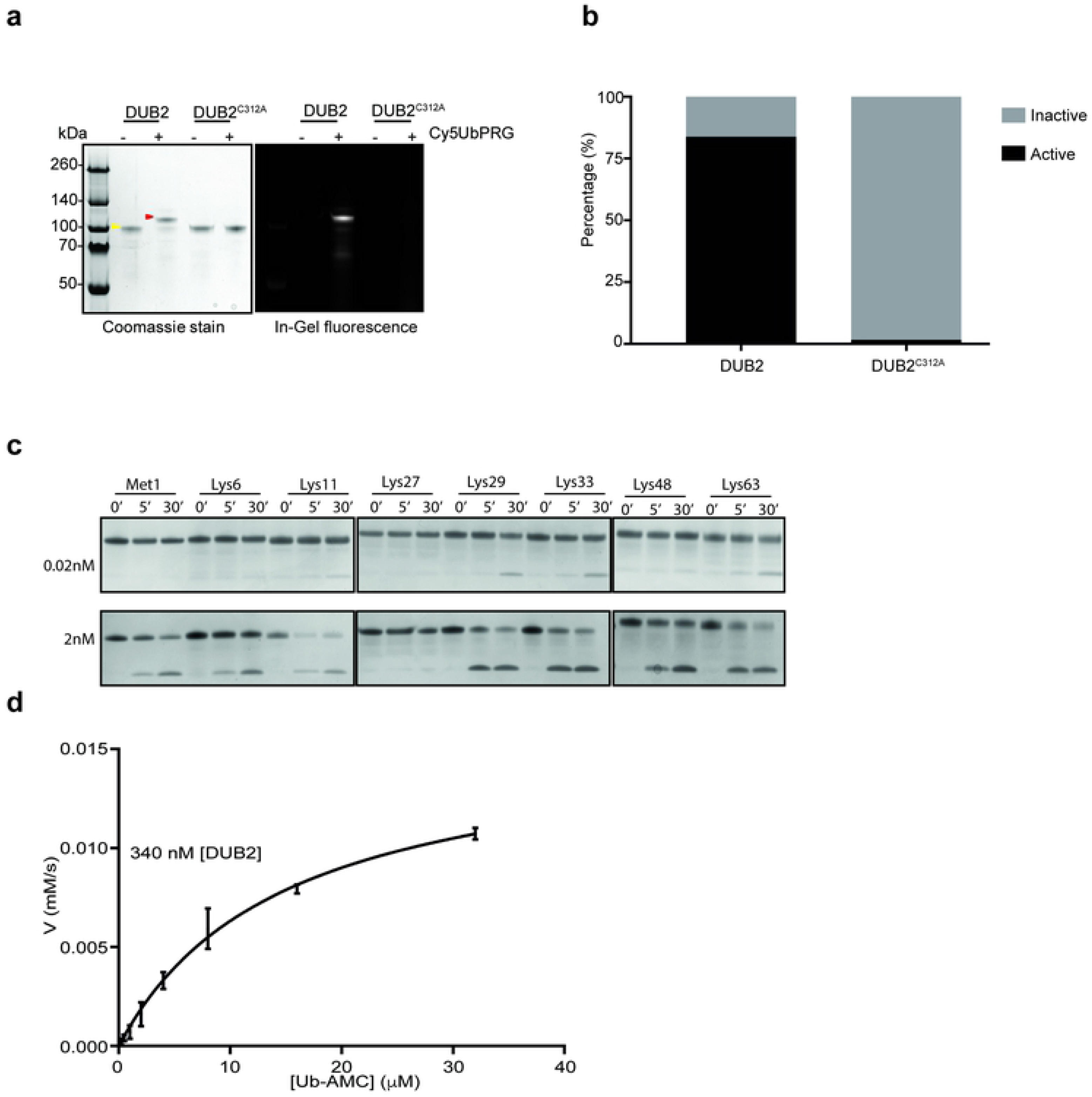
*In vitro* characterization of DUB2 recombinant protein. **a** 500 ng of purified DUB2 and DUB2^C312A^ were incubated with or without 1.2 μg of Cy5UbPRG for 30 min. Samples were analysed by SDS-PAGE gel electrophoresis with in-gel fluorescence. The gel was stained in InstantBlue^TM^ protein stain. Yellow arrowhead is DUB2 and the red arrowhead is DUB2 in complex with the Cy5UbPRG probe. **b** The stained gel was analysed using GelAnalyzer 19.1 software to determine the band intensity for DUB2 and DUB2 in complex with the ABP. Data were then collected and analysed to determine the ratio between active and inactive form of DUB2 in Prism software, where the graph was generated. **c** Different di-ubiquitin linkage types were incubated with recombinant DUB2 (0.02 nM or 2 nM) for the times indicated and proteins were resolved by SDS-PAGE gel and stained with InstantBlue^TM^ Protein Stain. **d** Michaelis-Menten kinetic analysis for DUB2 hydrolysis of Ub-AMC. Error bars represent s.d. from the mean. Reactions containing 400 nM DUB2 were initiated by the addition of Ub-AMC at final concentrations of 0.25 μM to 32 μM. Assays were performed in triplicate for three independent experiments. Each initial rate was determined by the gradient of a linear equation of the product concentration against the time (from 0-400 sec).

## Discussion

Recent studies showed the potential for proteasome inhibition as a treatment for three diseases caused by trypanosomatid parasitic protozoa, including leishmaniasis [9,10]. The proteasome is dependent on the ubiquitination pathway to identify substrates for degradation and a search of the *Leishmania* genome reveals a range of E1, E2 and E3 ligases, as well as a number of DUBs [33]. Nevertheless, there have been no extensive studies exploring the role and essentiality of *Leishmania* DUBs through the complex life cycle of the parasite. Our bioinformatic analysis identified 20 DUBs belonging to the C12, C19 and C65 peptidase families, suggesting an extensive deubiquitinating system in *L. mexicana*, comparable in size to yeast [34]. Additional cysteine peptidase DUBs from C78, C85 and clan CP C97 families have been characterised in another study [23]. Analysis of DUBs in promastigote lysates using the Cy5UbPRG ABP probe identified 6 active DUBs (DUB2, DUB15, DUB16, DUB17, DUB18 and DUB19) by mass spectrometry and additional active enzymes by in-gel fluorescence, demonstrating the presence of an active deubiquitinating system in *L. mexicana*. Additional active DUBs were identified by in-gel fluorescence comparison of wild type and gene deletion mutants (DUB6, DUB9, DUB13), yet many remain unidentified and could be active below the limit of detection or possibly expressed in life cycle stages in the sandfly that were not examined in this study. We also used the ABP to demonstrate that many of the DUBs have activity that appears to remain constant in the different life cycle stages, yet we also demonstrated that Δ*dub4*, Δ*dub7* and *Δdub13* had no detectable loss of fitness in the promastigote, but had a significant loss of fitness in the amastigote. Intracellular amastigotes have a stringent metabolic response that is acutely sensitive to metabolic perturbation [35]. Our data indicate that amastigotes are also highly sensitive to disruption of ubiquitination homeostasis.

The Bar-seq screen and individual viability assay of DUB null mutants highlighted DUBs with potential roles in differentiation and infection. Δ*dub4*, Δ*dub7* and *Δdub13* had the strongest loss of fitness defects observed in axenic amastigote, macrophage infection and mouse infection experiments. Additionally, Δ*dub3*, Δ*dub5*, Δ*dub6*, Δ*dub8*, Δ*dub10*, Δ*dub11* and Δ*dub14* showed a strong loss of fitness uniquely during mouse infection, perhaps due to increased selective pressures inside the host or the longer duration of the experiment. The identification of several DUBs essential for differentiation was not unexpected as the *Leishmania* proteome differs substantially between promastigote and amastigote stages [36]. DUBs can control the abundance of *Leishmania* proteins by rescuing them from proteasomal degradation or, more indirectly, by regulating autophagy. Autophagy has been shown to be essential for the differentiation of promastigotes into amastigotes both *in vitro* and *in vivo* [32,37]. Proteolysis is required for metacyclogenesis and the promastigote to amastigote transition [38] and an accumulation of ubiquitylated proteins in the cytosol has been documented at the onset of differentiation [39]. Therefore, a role for *Leishmania* DUBs in differentiation-associated protein degradation could explain the higher degree of essentiality observed for DUBs over other peptidases in our Bar-seq screen.

We were unable to generate null mutant lines for four DUBs, three C19 (DUB1, DUB2 and DUB12) and one C12 family (DUB16) peptidase. The successful generation of facilitated null mutant lines for the four DUBs provides a level of support that they are essential for the promastigote. DUB16 is related to human UCHL1 (33% identity) and UCHL3 (37% identity), for which one proposed role is the control of free ubiquitin levels through the processing of ubiquitin precursor and ubiquitinated proteins [40]. The primary cytoplasmic localisation of DUB16 supports the idea of a general role in regulating free ubiquitin levels, but the nuclear localisation of DUB12 points to a possible role in regulating fundamental nuclear processes such as transcription, RNA processing, DNA repair and/or epigenetic regulation. These processes have previously been linked to the activity of human DUBs [41]. USP36 is the human DUB most closely related to DUB12 (28% identity) and has been shown to regulate the stability of nuclear proteins including c-Myc, B23 and fibrillarin, the latter two of which are required for rRNA processing and ribosomal biogenesis [42]. USP36 deletion is lethal in mice, attributable to its role in rRNA processing and protein synthesis [43]. However, deletion of the yeast homologue, Ubp10, only results in a reduced growth phenotype [44]. In our DUB12 overexpression line, we observed an increase in the activity of two forms of DUB12, likely arising from its proteolytic cleavage to 55 kDa proteins from an 85 kDa precursor. The post-translation modification of DUBs is one way in which their activity has been shown to be regulated and, although rare, proteolytic cleavage of DUBs has been described for some human DUBs [45,46].

DUB2 is a cysteine peptidase with a C19-UCH domain, a UBA domain at the C-terminus and a ZnF_UBP domain at the N-terminus. The most similar human DUBs are USP5 and USP13. USP5 is a well characterised member of the USP family responsible for the disassembly of the majority of unanchored poly-ubiquitin *in vivo* [47]. The ZnF-UBP domain of USP5 recognises the C-terminus of ubiquitin [48] and USP5 has been shown to hydrolyse the Lys-63, Lys-48, Lys-11, Lys-29 linkages [49]. DUB2 is similar to USP5 in having a broad linkage specificity, towards all the di-ubiquitin chains except Met-1 and Lys-29 *in vitro*. Our *in vitro* experiments demonstrate that DUB2 exhibits deubiquitinating activity against both artificial DUB substrates (Cy5-Ub-PRG ABP and Ub-AMC) and di-ubiquitin. Several human DUBs have previously been demonstrated to have a pleotropic function by regulating protein stability or by regulating assembly and function of different machineries. DUB2 essentiality, localisation and interactome suggests a pleotropic role for DUB2 deubiquitination of multiple substrates. The possibility that DUB2 is responsible for the release of the unattached polyubiquitin chains could also explain the phenotype in *Leishmania* after DUB2 depletion. The sudden and significant depletion of free ubiquitin results in significant dysfunction of all molecular machineries associated with ubiquitin, such as proteasomal degradation, endocytosis, replication, transcription and RNA splicing. If this was the case the parasite would enter a ‘free ubiquitin crisis’, which would eventually lead to death.

The interactome data suggest DUB2 plays a role in post-transcriptional gene regulation. In *Leishmania* and other kinetoplastids gene expression occurs in an unusual manner including constitutive polycistronic transcription and *trans-*splicing of a splice leader RNA to generate mature mRNAs [50]. Amongst the DUB2 interactors, one of the most strongly enriched proteins was cyclin 12 which in trypanosomes forms a tripartite complex with CRK9 and a CRK9 associated protein, CRK9AP. The CYC12-CRK9-CRK9AP complex is essential for *trans*-splicing of the splice leader RNA [29]. However, CRK9 and CRK9AP were not detected in the DUB2 interactome, indicating that DUB2 plays a role in turnover of free CYC12. Depletion of CRK9AP causes a rapid loss of CYC12 at the protein but not mRNA level, suggesting dynamic regulation of CYC12 protein [29]. DUB2 may therefore protect free CYC12 from degradation, when DUB2 is absent, CYC12 may become a target for degradation leading to co-depletion of CRK9 and a critical loss of *trans* splicing capacity. Alternatively, deubiquitination of CYC12 may be required for complex formation, similar to the requirement for the removal of ubiquitin from Bcl10 by Usp9x for its tripartite complexing to Malt1 and Carma1 [51]. Another strongly enriched protein in the DUB2 interactome was the histone chaperone NAP1, an important factor in nucleosome assembly [30]. This implies a role for DUB2 in nucleosome dynamics. In other eukaryotes, ubiquitination/de-ubiquitination of H2A and H2B is a dynamic process that modulates the stability of nucleosomes to allow transcription/DNA replication to take place [52]. The transcription factors, splicing factors and DNA replication proteins co-enriched with DUB2 are consistent with a role in regulating nucleosome assembly/disassembly since this would place DUB2 in the vicinity of these processes.

DUB activity can be regulated by post-translational modifications such as phosphorylation, one of the best characterised examples is activation of OTUD5 by phosphorylation in the catalytic domain [53]. Ubiquitination can also affect DUB activity, for example ubiquitination of UCH-L1 near the active site prevents binding to ubiquitin [54]. We identified a phosphorylated region and a ubiquitination site flanking the DUB2 UBA domain which could similarly regulate binding of DUB2 to ubiquitinated substrates. DUB2 has a broad specificity for diubiquitin cleavage, with Lys-27 being the only linkage for which there is poor activity. Lys-11, Lys-29 and Lys-48 linked chains have roles in proteasomal degradation, Lys-29 is associated with lysosomal degradation and Lys-33 is involved in both secretory and endocytic pathways, acting as a sorting signal for membrane proteins [55]. Another preferred linkage of DUB2 is Lys-63 which can serve as a molecular glue, allowing the rapid and reversible formation of pivotal signalling complexes. Lys-63 ubiquitin chains have been shown to regulate several biological processes including DNA repair, clearance of damaged mitochondria, protein sorting and they can guide assembly of large protein complexes that drive mRNA splicing and translation [55]. In the DUB2 pull down, the VPS4 protein was identified, which could indicate a putative role of DUB2 in endosome sorting and could explain why DUB2 cleaves Lys-33 and Lys-29. Whilst VPS4 is the only known protein associated with autophagy/endocytosis that was identified in the DUB2 interactome, a role in the endocytic system fits with DUB2 lysosomal localisation. Previously, VPS4 was shown to play a role in endosome sorting and autophagy in *L. major*, which are essential for the differentiation and virulence of this parasite [32]. This might explain the upregulation of ConA uptake in DUB2-depleted cells.

In the past decade there has been an expanding interest in DUBs as potential drug targets as they have been shown to play a crucial role in both cancer and neurodegenerative diseases [56]. The determination of the structure of multiple DUBs, including catalytic domains, in combination with the development of advanced high-throughput screening-compatible assays with USP substrates has resulted in the discovery of several inhibitors against a select number of human DUBs including and USP7 [57,58] and USP14 [59]. *DUB2* is the first DUB to have been genetically validated as a drug target in *L. mexicana*. DiCre permitted the analysis of emerging phenotypes, confirming the essential role for DUB2 in promastigotes and in the establishment of mouse infection. Analysing *Leishmania* genes in this manner represents a high level of genetic validation [60]. To date, most drug target validation has relied on genetic manipulation of promastigotes, however, the Bar-seq approach presented here has revealed some DUBs to have amastigote-specific loss of fitness, making DUBs 3, 5, 6, 10, 11 and 14 potential drug targets. As DUB6 is related to human USP7 and DUB14 to human USP14, chemical tools developed for human DUB drug discovery programs [59] may be useful chemical entry points to develop inhibitors to the *Leishmania* enzymes.

## Materials and Methods

### Culture of Leishmania

*Leishmania mexicana* (MNYC/BZ/62/M379) were grown in HOMEM (Gibco) supplemented with 10% (v/v) heat-inactivated foetal calf serum (HIFCS) (Gibco) and 1% (v/v) Penicillin/Streptomycin solution (Sigma-Aldrich) at 25° C. Mid-log phase parasites were defined as between 4-8 × 10^6^ cells mL^−1^ and stationary phase parasites between 1.5-2.5 × 10^7^ cells mL^−1^.

*L. mexicana* promastigotes were differentiated to axenic amastigotes as previously described [61]. Briefly, stationary phase culture of *L. mexicana* was pelleted by centrifugation at 1,000 × g for 10 minutes and washed with 1 × PBS. Cells were then resuspended in amastigote culture medium (Schneider’s Drosophila medium [Gibco], 20% HIFCS and 0.015 mg mL^−1^ hemin [stock dissolved in 50 mM NaOH], pH 5.5) with a concentration of 2 × 10^6^ cells mL^−1^ in 6-well plates. Except where stated otherwise, cells were then incubated at 32°C with 5% CO2. For transformation of amastigotes back to promastigotes after 120 h, 100 μL of axenic amastigotes were diluted in 5 mL of HOMEM medium with 10% HIFCS and 1% Penicillin/Streptomycin and placed in 6-well plates. The plate was then incubated at 25°C. Transgenic parasites were grown with the following antibiotic concentrations: G418 (Neomycin) at 50 μg mL^−1^; Hygromycin at 50 μg mL^−1^; Blasticidin S at 10 μg mL^−1^; Puromycin at 30 μg mL^−1^ (antibiotics from InvivoGen). To assess viability assay of axenic amastigotes, 200 μL of axenic amastigotes at 1 × 10^6^ cells mL^−1^ were placed in 96 well plates and incubated at 35°C with 5% CO_2_. 20 μL of 0.05 μM resazurin (Sigma) was added and the plate was incubated overnight as before. The next day, resorufin fluorescence (579Ex/584Em) was measured using a Polarstar plate reader (Omega). To assess infection of mice, 10^6^ stationary phase *L. mexicana* promastigotes were injected into the right footpad of female BALB/c mice (Charles River Laboratories). Lesion size was monitored weekly. Parasite burden was assessed as previously described [62]. All experiments were conducted according to the Animals (Scientific Procedures) Act of 1986, United Kingdom, and had approval from the University of York Animal Welfare and Ethical Review Body (AWERB) committee.

*L. mexicana* transfections were performed using a T Cell Nucleofector™ Kit (Lonza) as described previously [23]. The transfected culture was split between two flasks to select for independent transfection events and incubated overnight at 25°C to recover. The next day, suitable antibiotics were added and cells cloned by serial dilutions of 1 in 6, 1 in 66 and 1 in 726 in 96-well microplates.

### Phylogenetic Analysis

The protein sequences for each DUBs were extracted from (Uniprot- https://www.uniprot.org/) [63]. For human DUBs the canonical sequence was used as it was determined by Uniprot. The protein alignment was performed using MUSCLE (multiple sequence alignment with high accuracy and high throughput). A constraint tree was created by including DUBs into the seven known families. Finally, a phylogenetic tree was generated by RAxML (https://raxml-ng.vital-it.ch/#/) [64]. The LG Substitution matrix was used. The best fit model tree was further designed initially in the iTOL INTERACTIVE TREE OF LIFE (https://itol.embl.de/) where branched length was ignored, and an unrooted tree style was formed. Finally, the Adobe Illustrator software was then used to finalise the tree.

### Life cycle phenotyping using Bar-seq

Null mutants were generated using a CRISPR-Cas9-based approach [21] with a cell line expressing Cas9 and T7 RNA polymerase [23]. DNA was harvested from clones or populations for diagnostic PCRs using the QIAGEN DNeasy Blood and Tissue Kit. Two sets of primers were designed to confirm the generation of null mutants, the first specific to the ORF of the target gene and the second specific to the UTRs (Forward Primer 5’-UTR, Reverse Primer 3’-UTR). Null mutant lines were grown to mid-log phase, spun down at 1,000 × g for 10 min and pooled in equal proportions in Grace’s medium (Grace’s insect media [Sigma-Aldrich] with 4.2 mM NaHCO_3_, 10% FCS [Gibco], 1 × Penicillin/Streptomycin [Gibco] and 1 × BME Vitamins [Sigma-Aldrich] and adjusted to pH 5.25) up to a total concentration of 2 × 10^6^ cells mL^−1^. Cultures were prepared in sextuplicate to provide technical replication. Next, the pooled cells were grown at 25°C for 7 d to allow for the enrichment of metacyclic promastigotes. To prepare axenic amastigotes from the enriched metacyclic population, cells were spun down and resuspended in amastigote medium at 2 × 10^6^ cells per well. Cells were then cultured at 35°C with 5% CO_2_.

Bone marrow-derived macrophages were extracted from female BALB/C mouse femur, equilibrated in warm DMEM (GE Life Sciences), spun at 200 × g for 10 min and resuspended in macrophage medium (DMEM [Invitrogen] plus 10% FCS and 2 mM L-glutamine) in order to have 2.5 × 10^6^ cells per well. Following overnight incubation at 37°C, enriched metacyclic cells were purified using a Ficoll 400 (Sigma-Aldrich) gradient (40%, 10% and 5% Ficoll) and centrifuged at 1,300 × g for 10 min. Following washing with Grace’s medium, metacyclic promastigotes were added to the macrophages in a 1:1 ratio and left to interact for 4 h at 35°C with 5 % CO_2_. Excess *Leishmania* cells were then removed by washing with DMEM and the cells incubated in DMEM 10% HIFCS for 12 h and 72 h.

To perform the mouse infections, 10^6^ purified metacyclic promastigotes were injected into the left footpad of BALB/c mice. Mice were culled at 3 and 6 weeks post-inoculation. At the relevant time points, DNA was extracted from experimental samples and processed using the QIAGEN DNeasy Blood and Tissue Kit. DNA samples were prepared for next-generation sequencing by first amplifying unique barcodes by PCR using the following primers: 5’-TCGTCGGCAGCGTCAGATGTGTATAAGAGACAGAgatgatgattacTAATACGACTCACTATAAAACTGGAAG-3’ and 5’-GTCTCGTGGGCTCGGAGATGTGTATAAGAGACAGGAGAGACAGGCATGCCTT-3’ containing the Illumina adapter sequence. Reactions were set up with Q5® polymerase (NEB) as per the manufacturer’s instructions except for the following changes: just 0.5 x of Q5 Reaction Buffer and Q5 High GC Enhancer were used. The cycling conditions were 98°C for 5 min followed by 20-25 cycles of 98°C for 30 sec, 60°C for 30 sec and 72°C for 10 sec. Following purification of the PCR reactions using the QIAGEN MinElute PCR Purification Kit, a second PCR was performed using Illumina’s Nextera XT indexing primers in order to add unique barcode sequences to each sample. 8 cycles of PCR amplification were performed using NEBNext® Q5 Polymerase 2X Master Mix (New England Biolabs), according to the manufacturer’s guidelines. Indexed amplicons were purified using 0.9 X AMPure XP beads (Beckmann Coulter) and eluted into low TE buffer before quantitation and pooling at approximately equimolar ratios. Amplicon pools were then sent for 150 base paired end sequencing on an Illumina HiSeq 3000 sequencer at the University of Leeds Next Generation Sequencing Facility.

Illumina read sequences were analysed using a custom Python script to search for the 12 bp sequence preceding the barcode. If an exact match was found, the following 12 bp sequence was counted as barcode sequence. A total count for each unique barcode sequence was generated for each sample. Next, the relative fitness of null mutants at each life cycle stage was calculated by taking the number of unique barcode reads for each null mutant line and dividing it by the total number of expected barcode reads. Significant increases or decreases in null mutant line fitness between adjacent samples in the experimental workflow were analysed using paired t-tests and the Holm-Sidak method in GraphPad Prism 7.

### Construction of plasmids

A full list and descriptions of all primers and plasmids used in this study can be found in Table S3. The construction of the plasmids needed for the generation of *DUB2* inducible gene deletion was performed as previously described [27]. The *DUB2* gene including stop codon was cloned into pGL2315, creating pGL2727. For the complementation plasmids, pNUS-*DUB1, DUB2, DUB12 and DUB16* expression plasmids were generated using Gibson Assembly. The successful clones were confirmed using PCR and sequencing.

### Induction of diCre mediated gene deletion

DiCre in cell lines containing both diCre and loxP flanked genes of interest was induced by the addition of between 100 nM and 1 μM rapamycin (Abcam). Promastigotes were then allowed to grow for 48 h and split into new cultures with a concentration of 1 × 10^5^ cells mL^−1^ and induced again with 100 nM of rapamycin.

### Preparation of protein extracts

3 × 10^7^ *L. mexicana* cells in the desired life cycle stage were pelleted by centrifugation at 1,000 g for 10 min. Cells were washed in 1 x PBS and lysed by resuspending in 40 μL of 1 x SDS-PAGE loading buffer. Next, samples were heated for 10 minutes at 90°C on a heat block. Samples were allowed to cool down at room temperature and 250U of Expedeon BaseMuncher endonuclease was added to each sample. Samples were then incubated at 37°C for 30 minutes and loaded directly into a NuPAGE 4-12% Bis-Tris protein gel (Thermofisher) or stored at −20° C.

### Cy5UbPRG profiling

3 × 10^7^ cells from a log-phase *L. mexicana* culture or 9 × 10^7^ amastigote were spun at 1,000 × g and washed three times with 1 mL of ice-cold TSB Wash buffer (44 mM NaCl, 5 mM KCl, 3 mM NaH2PO4, 118 mM sucrose and 10 mM glucose, pH 7.4). Next, the cells were lysed using a newly prepared ice-cold lysis buffer (50 mM Tris-HCl pH 7.4, 120 mM NaCl, 1% NP40 and freshly added in order: 1 μg mL^−1^ pepstatin, 1 x cOmplete™ ULTRA Tablets, [Mini, EASYpack Protease Inhibitor Cocktail, Roche], 1 mM DTT, 1 mM PMSF, 0.01 mM E64). Samples were incubated at 4°C for 15 minutes. Afterwards, samples were centrifuged at 17,000 × g for 15 minutes and the supernatant withdrawn. Samples were prepared to have a protein concentration of 1 mg mL^−1^ in a total volume of 25 μL. 2 μL of 50 mM NaOH and 1 μM of Cy5UbPRG (UbiQ) were added. Lysis buffer was used to top up to a final volume of 25 μL. The reaction was then incubated at room temperature for 30 min and stopped by the addition of 3 x loading dye. 12 μL were then analysed in a gradient 4-12% NuPage Bis-Tris protein gel, for exactly 90 min at 200 V. The gel then was washed with water and imaged with Amersham Typhoon with Excitation: 635 nm, Filter: Cy5 670BP30 (GE Healthcare Life Sciences).

For purification of DUBs cyanogen bromide (CNBr)-activated Sepharose 4B resin (GE Healthcare) was used to couple with Ub-PRG (1.2 pmol) or Ubiquitin (1.2 pmol) according to the manufacturer’s protocol (GE Healthcare). Beads were then stored in 4°C until used. 50 mg of the coupled resin of either Ub-PRG or ubiquitin were incubated with fresh *Leishmania* lysate collected from 1 x10^8^ cells (prepared as described above) at 37°C for 3 h. The beads were then washed with PBS plus 1% Triton X-100 (3 × 10 mL), 2% CHAPS, 8 M Urea in PBS and distilled water. The bound proteins were then digested on-bead with trypsin and analysed by MS/MS spectrometry. The MS data was processed using Data Analysis software (Bruker) and the automated Matrix Science Mascot Daemon server (v2.4.1). Protein identifications were assigned using the Mascot search engine to interrogate protein sequences in the annotated proteins database (obtained from TriTrypDB), allowing a mass tolerance of 0.4 Da for both MS and MS/MS analyses.

### Immunoprecipitation of myc-tagged *Leishmania* proteins

1 × 10^9^ parasites were washed twice in PBS and cross-linked with 1 mM dithiobis(succinimidyl propionate) (DSP, Thermo Scientific) in 10 ml PBS for 10 min at 26°C. DSP was quenched with 20 mM Tris pH 7.5 for 5 min. Parasites were lysed in ice cold 500 μl lysis buffer (1% NP40, 50 mM Tris pH 7.5, 250 mM NaCl, 1 mM EDTA, 0.1 mM PMSF, 1 μg mL^−1^ pepstatin A, 1 μM E64, 0.4 mM 1-10 phenanthroline, 20 μl mL^−1^ proteoloc (Expedeon), 0.17 complete protease inhibitor tablets mL^−1^ (Sigma)) by probe sonication for 3 × 5 sec. Lysate was centrifuged for 10 min at 10,000 × g at 4°C and protein concentration in the supernatant measured by BCA assay (Thermo scientific). Lysate equivalent to 5 mg of total protein was incubated with 30 μl anti-myc magnetic beads (Thermo scientific) for 2.5 hr at 4°C with rotation. Beads were washed 4 × 300 μl ice cold lysis buffer for 5 min each wash followed by two PBS washes. Beads were then stored at −80°C. Myc-tagged protein was eluted at room temperature with 25 μl myc-peptide (0.5mg/mL in PBS, Sigma) for 15 minutes at 700 rpm mixing. The elution step was repeated and eluates pooled. 1/10^th^ of the elution was used to check successful immunoprecipitation by western blot. The remainder was mixed with 4 volumes of absolute methanol and 1 volume of chloroform and vortexed for 1 min. The sample was then centrifuged for 1 h at 18 000g at 4°C. The pellet was washed with 270 μL of absolute methanol and centrifuged for 10 min at 18,000 × g at room temperature. The pellet was resuspended in 150 μL 50 mM TEAB pH 8.5, 0.1% PPS silent surfactant (Expedeon) for 1 h by shaking at 800 rpm at room temperature. Afterwards, 10 mM of Tris (2-carboxyethyl) phosphine (TCEP) and 10 mM Iodoacetamide (IAA) were added to the sample and incubated for 30 min at room temperature in the dark. Finally, 200 ng of trypsin and 1 mM CaCl_2_ were added and proteins digested overnight at 37°C 200 rpm. After digestion, PPS silent surfactant was cleaved by acidifying the digest to 0.5% trifluoroacetic acid (TFA) and incubating 1 h at RT. Digest was centrifuged for 10 min at 17,000 × g. Peptides were desalted with C18 (3M Empore) desalting tips.

Samples were loaded onto an UltiMate 3000 RSLCnano HPLC system (Thermo) equipped with a PepMap 100 Å C18, 5 µm trap column (300 µm × 5 mm Thermo) and a PepMap, 2 µm, 100 Å, C18 EasyNano nanocapillary column (75 μm × 150 mm, Thermo). The trap wash solvent was aqueous 0.05% (v:v) trifluoroacetic acid and the trapping flow rate was 15 µL min^−1^. The trap was washed for 3 min before switching flow to the capillary column. Separation used gradient elution of two solvents: solvent A, aqueous 1% (v:v) formic acid; solvent B, aqueous 80% (v:v) acetonitrile containing 1% (v:v) formic acid. The flow rate for the capillary column was 300 nL min^−1^ and the column temperature was 40°C. The linear multi-step gradient profile was: 3-10% B over 7 min, 10-35% B over 30 min, 35-99% B over 5 min and then proceeding to wash with 99% solvent B for 4 min. The column was returned to initial conditions and re-equilibrated for 15 min before subsequent injections.

The nanoLC system was interfaced with an Orbitrap Fusion Tribrid mass spectrometer (Thermo) with an EasyNano ionisation source (Thermo). Positive ESI-MS and MS2 spectra were acquired using Xcalibur software (version 4.0, Thermo). Instrument source settings were: ion spray voltage, 1,900 V; sweep gas, 0 Arb; ion transfer tube temperature; 275°C. MS1 spectra were acquired in the Orbitrap with: 120,000 resolution, scan range: m/z 375-1,500; AGC target, 4e5; max fill time, 100 ms. Data dependent acquisition was performed in top speed mode using a 1 sec cycle, selecting the most intense precursors with charge states >1. Easy-IC was used for internal calibration. Dynamic exclusion was performed for 50 sec post precursor selection and a minimum threshold for fragmentation was set at 5e3. MS2 spectra were acquired in the linear ion trap with: scan rate, turbo; quadrupole isolation, 1.6 m/z; activation type, HCD; activation energy: 32%; AGC target, 5e3; first mass, 110 m/z; max fill time, 100 msec. Acquisitions were arranged by Xcalibur to inject ions for all available parallelizable time.

Peak lists in .raw format were imported into Progenesis QI (Version 2.2., Waters) and LC-MS chromatograms aligned. Precursor ion intensities were normalised against total intensity for each acquisition. A combined peak list was exported in .mgf format for database searching against the *L. mexicana* subset of the TriTrypDB database (8,250 sequences; 5,180,224 residues). Mascot Daemon (version 2.6.1, Matrix Science) was used to submit the search to a locally-running copy of the Mascot program (Matrix Science Ltd., version 2.6.1). Search criteria specified: Enzyme, trypsin; Max missed cleavages, 2; Fixed modifications, Carbamidomethyl (C); Variable modifications, Oxidation (M), Phosphorylation (S,T,Y); Peptide tolerance, 3 ppm; MS/MS tolerance, 0.5 Da; Instrument, ESI-TRAP. Peptide identifications were passed through the percolator algorithm to achieve a 1% false discovery rate as assessed against a reversed database and individual matches further filtered to minimum expect scores of 0.05. The Mascot .XML result file was imported into Progenesis QI and peptide identifications associated with precursor peak areas. Relative protein abundance was calculated using precursor ion areas from non-conflicting unique peptides. Accepted protein quantifications were required to contain a minimum of two unique peptide matches. Interactors were scored using SAINTq [65], using a false discovery rate threshold of <1% to select high confidence DUB2 interactors.

### Protein expression and purification

A baculovirus expression system was used for the expression of DUB2 recombinant protein in sf9 insect cells as described in the Bac-to-Bac Baculovirus Expression protocol (Invitrogen). The gene was first cloned into the PFastBacNKI-his-3C-LIC vector using Ligation Independent Cloning (LIC). After three days, the insect cells were lysed in lysis buffer (30 mM Tris-HCl, pH 8, 0.3 M NaCl, 0.03 M imidazole, 5 mM β-mercaptoethanol and 1 tablet of cOmplete™ ULTRA Tablets, Mini, EASYpack Protease Inhibitor Cocktail, Roche) by sonication. The samples were then centrifuged (21 K, 4°C, 30 min) and the supernatant applied to an AKTA Start (GE Life Sciences) at 5 mL min^−1^, to a 5 mL His-trap crude FF column (GE Life Sciences). This was washed with Buffer A (30 mM Tris-HCl pH. 8, 0.3 M NaCl, 0.03 M imidazole, 5 mM β-mercaptoethanol) and the protein eluted with increasing concentration of imidazole (0.03-0.5 M). Protein was concentrated using Amicon Ultra-4 30k MWCO (Millipore). 3C protease was added to purified protein in a 1:50 (w:w, 3C:Recombinant protein) ratio with 2 mM DTT. The aliquot was added to a 16 kDa dialysis tube and incubated at 4°C in buffer A overnight. Protein was loaded onto a 5 mL His-trap crude FF column and ran as described above. The flow-through from the His-tag cleavage containing the tag free protein was concentrated to 2 mL using Amicon Ultra-4 30k MWCO (Millipore). The concentrated protein was applied to a HiLoad 16/600 S75pg column (GE Life Sciences), equilibrated and eluted in 25 mM Tris-HCl pH 8.0, 150 mM NaCl, 5 mM β-mercaptoethanol. Fractions containing the protein were identified by absorbance at 280 nm and analysed by 10% SDS-PAGE. These fractions were pooled and concentrated up to 2 mg mL^−1^. 1 mM of DTT was added to the sample and the protein stored at −80°C.

### Ubiquitin Probe Assay for Recombinant protein

10 μL of the appropriate concentration of the recombinant DUB was incubated with 2 μL 50 mM NaOH, 5 μL dH_2_O and 2 μL 0.25 μg mL^−1^ ubiquitin probe Cy5UbPRG (UbiQ) for 30 min at room temperature. Binding of probe was analysed using a NUPAGE™ 4-12% Bis-Tris Gel. The gel was then scanned using the Amersham Typhoon (GE Healthcare Life Sciences) and stained with InstantBlue^™^ Protein Stain.

The in *vitro* enzymatic assay with the fluorogenic substrate ubiquitin-7-amido-4-methylcoumarin (Ub-AMC, UbiQ) was assembled in a 384-well microplate (Thermo Scientific). Assays were performed in 20 μL reaction volumes and in triplicate. The reactions were initiated by the addition of 10 μL of substrate or enzyme. Substrate and enzymes were diluted in buffer containing 50 nM Tris, pH 8.0, 0.15 M NaCl, 5 nM DTT and 0.05% CHAPS. The fluorescence intensity was monitored with a Polarstar plate reader (Omega) equilibrated at 25°C at intervals of 20 sec for 20 min using excitation/emission filter pairs of 340/460 nm. The fluorescence value was then converted into concentration using a standard curve. The standard curve was determined by allowing the reactions to plateau and the maximum arbitrary fluorescence units were measured and plotted against a known substrate concentration. An AMC concentration against time graph was generated in Excel software and the initial slope of each reaction was determined by generating a linear regression. For the determination of the Km and kcat values of DUB2, the enzyme concentrations were kept constant, and the concentration of Ub-AMC varied from 0.25 μM to 32 μM. The initial rates in μM sec ^−1^ were then plotted against substrate concentration (μM) in Prism 7 software. The data were fitted to the Michaelis-Menton equation so the kinetic parameters, Km and kcat were determined.

## Acknowledgments

This work was supported by Medical Research Council studentships to AD and RB (MRC MR/N018230/1). Additional support was provided by the MRC (MR/K019384/1) and Wellcome Trust (200807/Z/16/Z). We would like to thank UbiQ for supplying Ub-AMC and Cy5UbPRG.

## Supplementary Data

**Fig. S1: The generation of the DUB null mutant library**

Diagnostic PCRs that were performed to check the successful generation of null and facilitated null mutants. PCR analysis of genomic DNA of at least two individual clones (null mutants, CL1, CL2 etc) or two separate populations (facilitated null mutant, **c**), using the indicated set of primers (ORF specific primers **(a)**, UTR’ specific primers **(b)**). As a control, the parental T7 Cas9 cell line was used. The resulting amplicons were resolved on a 1% agarose gel and stained with SYBR safe. The expected size of the resulting amplicons is presented in Table S3. Schematic representation of the wild type, heterozygous, null mutant and facilitated null mutant genomic loci including the diagnostic primers (green arrows primers bind to ORF, grey arrow primers bind to UTRs and amplify across the loci, purple arrow binds to the blasticidin resistance marker).

**Fig. S2: Activity profiling of the DUB null mutant cell lines**

**a** Lysate extracted from log-phase *L. mexicana* promastigotes treated with or without Cy5UbPRG for 30 min. Proteins were separated by SDS-PAGE and in-gel fluorescence was captured using a Typhoon imager followed by Coomassie staining as a loading control. **b** Lysates extracted from null mutant lines of a log-phase *L. mexicana* promastigotes treated with Cy5UbPRG for 30 min. In-gel fluorescent images were obtained as under a. The red arrowhead shows the position where an active DUB is missing compared to the parental T7 Cas9 cell line. **c** mNeonGreen (mNeon) endogenously N-terminally tagged DUB lines (mNeon:DUB) were assayed using the Cy5UbPRG activity probe. Samples were lysed and incubated for 30 min with Cy5UbPRG probe. In-gel fluorescent images were obtained as under a. Red represents a decrease in band intensity (the endogenous DUB), whereas blue represents an increase (DUB:mNeon).

**Fig. S3: Expression of HASPB in fully differentiated amastigotes**

**a** Western blot analysis of 2 × 10^7^ axenic amastigotes. Samples were separated in a 4-15% protein gel. The stain-free gel used contains trihalo compounds which in the presence of the UV-light react with tryptophan residues, producing fluorescence. The gel was activated by 45 sec UV exposure, proteins were transferred to a PVDF membrane and probed with 1:1,500 dilution of anti-HASPB. Finally, as a loading control the total protein was determined using the stain-free property of the gel. **b** Lysate extracted from differentiated promastigotes to amastigotes (48 h and 144 h after initiation of axenic differentiation) treated with or without Cy5UbPRG for 30 min. Protein was separated in an SDS-gel, and the image was captured using Typhoon and then stained with Coomassie. **c** Western blot analysis of 2 × 10^7^ axenic amastigote grown for 144 h, as above.

**Fig.S4: Localisation of endogenous tagged *Leishmania* DUBs**

Imaging of parasites using confocal microscopy and processed in ZEN 2.3 software. Panels: Hoechst 33342 for staining DNA (blue); top right, the green fluorescence signal from mNeon green; bottom left, Differential Interference Contrast (DIC); bottom right, merged.

**Fig. S5: Generation and validation of *DUB2* inducible cell line**

**a** Left: PCR analysis of extracted gDNA demonstrates successful integration of *HYG* cassette. The replacement of *DUB2* wild type allele with the *HYG* cassette was detected by PCR amplification using the primers shown in the schematic (Right). The forward primer was designed to bind to the ORF of *DUB2* whereas the reverse primer binds on the 3’ UTR of the target gene. Black arrows represent the primers.

**b** PCR amplification of *DUB2*^−/+FLOX^ Clone 4, 6, *DUB2*^−/+FLOX^ [*DUB2*] and *DUB2*^−/+FLOX^ [*DUB2^C312A^*] surviving parasites after treatment with RAP in the clonal assay (Fig.4e). Schematic representation of the primers used for the PCR as well as the expected fragments are shown in Figure 4A.

**c** Flow cytometry strategy to detect ConA uptake and cell size after RAP treatment in *DUB2*^+/+FLOX^ and *DUB2*^−/+FLOX^ clone 4 and clone 6 promastigotes. Promastigotes were grown in the presence or absence of 100 nM RAP for 48 h. Afterwards, the cells were seeded at a density of 1 × 10^5^ cells mL^−1^ and allowed to grow in the presence and absence of 100 nM RAP for another 24 h. Forward scatter was used to determine the size of the cells and FITC to determine ConA uptake. Red represents the non-induced promastigote population whereas blue represents the induced cell population.

**c** Compiled flow cytometry results of the forward scatter (Left) and ConA uptake (Right) in DUB2^+/+flox^ and DUB2^−/+flox^ promastigotes after 24 h post induction. Cells were grown for 48 h with or without RAP. Afterwards, the cells were seeded at a density of 1 × 10^5^ cells mL−1 and allowed to grow in the presence or absence of 100 nM of RAP for 24 h. The data were analysed by FlowJo_v10, where the mean of forward scatter was determined. Finally, the figure was generated in Prism Software. Each band represents a mean of 2 biological replicates. Error bars represent SEM.

**Fig. S6: DUB2 Immunoprecipitation using anti-MYC beads.**

**a** DUB profiling using the ABP Cy5UbPRG. Cell extract from a WT and mNeon:DUB2 was incubated with Cy5UbPRG for 30 min. Proteins were then separated by SDS-PAGE and imaged. The yellow arrow represents the DUB2 WT band whereas the red arrow represents the mNeon-DUB2. **b** Western blotting analysis with anti-MYC antibody (1:2,000) of protein cell extract from samples collected after immunoprecipitation of control cell line (WT) and mNeon:DUB2. E1, E2 were samples collected from the elution (myc peptide 0.5 mg mL^−1^ in PBS).

**Fig. S7: Purification of DUB2 active (Left) and DUB2^C256^ inactive protein (Right).**

**a** Protein gel of the soluble fraction (B), flow though (FT) and eluted fractions across the peak that were collected after Ni^2+^ affinity chromatography. 10 μL of the protein samples were separated by SDS-PAGE and visualised by InstantBlue^TM^ Protein Stain. **b** Protein gel of the sample before overnight dialysis with protease 3C for the removal of the his-tag (B), after overnight dialysis (A) and flow through and eluted fractions across the peaks collected after the second affinity chromatography. **c** Protein gel of the diluted (D) and concentrated (C) samples before application to a HiLoad 16/600 S75pg column and eluted fractions collected after size-exclusion chromatography.

**Table S1:** *Leishmania mexicana* C12, C19 and C65 deubiquitinases.

**Table S2:** DUB2 interacting partners

**Table S3:** Oligonucleotide primers

**Table S4:** Bar-seq data

**Table S5:** DUB2 interacting partner data

**Table S6:** DUB phosphorylation sites

